# Assessing the stability of polio eradication after the withdrawal of oral polio vaccine

**DOI:** 10.1101/084012

**Authors:** Michael Famulare, Christian Selinger, Kevin A. McCarthy, Philip A. Eckhoff, Guillaume Chabot-Couture

## Abstract

The oral polio vaccine (OPV) contains live-attenuated polioviruses that induce immunity by causing low virulence infections in vaccine recipients and their close contacts. Widespread immunization with OPV has reduced the annual global burden of paralytic poliomyelitis by a factor of ten thousand or more and has driven wild poliovirus (WPV) to the brink of eradication. However, in instances that have so far been rare, OPV can paralyze vaccine recipients and generate vaccine-derived polio outbreaks. To complete polio eradication, OPV use should eventually cease, but doing so will leave a growing population fully susceptible to infection. If poliovirus is reintroduced after OPV cessation, under what conditions will OPV vaccination be required to interrupt transmission? Can conditions exist where OPV and WPV reintroduction present similar risks of transmission? To answer these questions, we built a multiscale mathematical model of infection and transmission calibrated to data from clinical trials and field epidemiology studies. At the within-host level, the model describes the effects of vaccination and waning immunity on shedding and oral susceptibility to infection. At the between-host level, the model emulates the interaction of shedding and oral susceptibility with sanitation and person-to-person contact patterns to determine the transmission rate in communities. Our results show that inactivated polio vaccine is sufficient to prevent outbreaks in low transmission rate settings, and that OPV can be reintroduced and withdrawn as needed in moderate transmission rate settings. However, in high transmission rate settings, the conditions that support vaccine-derived outbreaks have only been rare because population immunity has been high. Absent population immunity, the Sabin strains from OPV will be nearly as capable of causing outbreaks as WPV. If post-cessation outbreak responses are followed by new vaccine-derived outbreaks, strategies to restore population immunity will be required to ensure the stability of polio eradication.

**Author Summary:** Oral polio vaccine (OPV) has played an essential role in the elimination of wild poliovirus (WPV). OPV contains attenuated yet transmissible viruses that can spread from person-to-person. When OPV transmission persists uninterrupted, vaccine-derived outbreaks occur. After OPV is no longer used in routine immunization, as with the cessation of type 2 OPV in 2016, population immunity will decline. A key question is how this affects the potential of OPV viruses to spread within and across communities. To address this, we examined the roles of immunity, sanitation, and social contact in limiting OPV transmission. Our results derive from an extensive review and synthesis of vaccine trial data and community epidemiological studies. Shedding, oral susceptibility to infection, and transmission data are analyzed to systematically explain and model observations of WPV and OPV circulation. We show that in high transmission rate settings, falling population immunity after OPV cessation will lead to conditions where OPV and WPV are similarly capable of causing outbreaks, and that this conclusion is compatible with the known safety of OPV prior to global cessation. Novel strategies will be required to ensure the stability of polio eradication for all time.

## Introduction

Wild polioviruses (WPV) have been eliminated from all but three countries [1, 2] by mass vaccination with the oral polio vaccine (OPV). The annual burden of paralytic polio infections has been reduced ten-thousand-fold since the start of vaccination efforts [1]. OPV has been the preferred vaccine for polio eradication because it costs less, can be reliably delivered by volunteers without medical training, and is more effective against poliovirus infection, relative to the inactivated polio vaccine (IPV) [3, 4]. Unique among current human vaccines, the live-attenuated Sabin poliovirus strains in OPV are transmissible. This transmissibility provides additional passive immunization that enhances the effectiveness of OPV for generating herd immunity. However, the attenuation of Sabin OPV is unstable and so it can, in rare instances, cause paralytic poliomyelitis [5] and lead to outbreaks of circulating vaccine-derived poliovirus (cVDPV) with virulence and transmissibility comparable to that of wild poliovirus (WPV) strains [6]. Thus, to complete the task of poliovirus eradication, vaccination with Sabin OPV must eventually cease [7].

The dual role of Sabin OPV as both a vaccine and a source of poliovirus is responsible for key uncertainties surrounding the ability of the Global Polio Eradication Initiative to achieve and sustain poliovirus eradication. Since the widespread introduction of polio vaccination, polio outbreaks have taken place in regions of low immunity against infection surrounded by regions of high immunity [8], OPV campaigns implemented in outbreak response have been effective for interrupting transmission [3], and cVDPV outbreaks have been rare consequences of the hundreds of millions of OPV doses administered every year [9].

However, after vaccination with OPV is stopped, population immunity against infection will progressively decline. If polioviruses are reintroduced, whether due to accidental or deliberate release [10–12], or because of sustained yet undetected transmission [2, 13–15], then large outbreaks may again occur. Outbreak control would require vaccination campaigns in affected countries and perhaps much more broadly, as has been done following recent type 2 cVDPV detections in Nigeria, Pakistan, the Democratic Republic of Congo and Syria [16–19].

In this paper, we explore the implications of the accumulated evidence about polio infection and transmission for the long-term stability of polio eradication. Will it be possible to interrupt all polio outbreaks without restarting widespread OPV vaccination, now and at all times in the future? Fundamental to this question is the ability of the Sabin polioviruses to circulate in low immunity populations. After OPV cessation, under what conditions will Sabin OPV remain the most effective tool for eliminating outbreaks without significant risks of causing more?

To address these questions, and building from primary literature and previous reviews and models [4, 20–25], we developed a comprehensive synthesis of the evidence for how within-host immunity, viral infectivity, and transmission dynamics fit together to explain the epidemiology of poliovirus transmission. We built a within-host model that summarizes the effects of immunization on poliovirus shedding and susceptibility. We then incorporated the within-host dynamics into a poliovirus transmission model using a household–community network framework, and we calibrated the model to field transmission studies. With the model, we explored how the average transmission rate in exposed communities varies with immunity, sanitation, number of social contacts, and poliovirus type. We identified conditions required for the Sabin strains to remain indefinitely as highly effective vaccines with low risks of causing outbreaks, and conditions where they can be expected to transmit nearly as efficiently as WPV. Our results are discussed in the context of the established stability of OPV cessation in the developed world and the ongoing global Sabin 2 cessation.

Previous models have also explored the effects of OPV cessation on Sabin and WPV transmission [20, 26–33]. Our work shares a similar emphasis on individual immunity and infection dynamics [20, 24, 34], but it differs in model structure, use of transmission data sources, and approach to epidemiological inference. The earlier work used compartmental models that assume individuals in large populations interact randomly [20, 26, 27, 33]. Models of specific settings—places and times—were based on national polio surveillance data [27, 33], and poliovirus evolution was modeled by extrapolation between Sabin and WPV endpoints based on assumed intermediate transitions [26, 27, 33]. In contrast, our model is based on person-to-person transmission among family members and extra-familial contacts. We calibrate specific settings to contact transmission data from field studies designed for that purpose [35–37]. We explore the effects of evolution by focusing on differences between the Sabin and WPV endpoints, and not the largely unstudied population genetics in between. In short, our model takes a bottom-up approach to modeling poliovirus transmission that complements existing work. Instead of drawing inferences about the unobserved conditions that affect transmission from observed outbreaks [27, 33], we draw inferences about unobserved properties of possible outbreaks from observed conditions that directly affect transmission.

## Methods

### Overview

To integrate knowledge of within-host immunity, shedding, and acquisition with between-host transmission, we built a multi-scale mathematical model. We first performed a quantitative literature review of clinical trial data to determine the impact of polio vaccination schedules containing OPV and/or IPV on poliovirus shedding after challenge with OPV. This resulted in an indirect measure of immunity—the OPV-equivalent antibody titer—which was used to model the associations between polio vaccination and shedding duration, concentration of virus in stool, and oral susceptibility to infection. Second, we reviewed in detail three historical transmission studies to parameterize a model of poliovirus transmission within households and between close extra-familial contacts. We then extended the person-to-person model by defining the local reproduction number—a threshold parameter that summarizes the potential for epidemic transmission within a community that has homogeneous demographics, immune histories, and sanitation practices. The model was implemented in Matlab 2015b (The Mathworks, Natick, MA) and is available in S1 Structured Code and Data and at famulare.github.io/cessationStability/. For all model parameters, see S2 Text.

We made simplifying assumptions while developing the model. First, we did not include the oral-oral transmission route. The studies known to us show that oral shedding occurs for shorter durations than fecal shedding and most often in individuals with low immunity [38–41], and this route is likely more important in high sanitation settings [34, 39, 42]. Second, we ignored fascinating questions about the effects of genetic evolution on Sabin strain transmission, and so our Sabin transmission parameters are most applicable in the first few weeks after OPV vaccination [43–45]. Third, our model focuses on transmission from children to family members and extrafamilial contacts, and it ignores other person-to-person interactions and possible environmental transmission routes, all of which influence the absolute probability and severity of outbreaks [46–50]. In the Results, we explore how the limitations of a model with these assumptions are informative about the roles of transmission route, viral evolution, and contact structure in various settings. Fourth, since paralysis has no direct influence on transmission, we did not model the impact of vaccination on paralysis (see Vidor *et al* [51] for a review).

All model features that describe the fraction of subjects shedding after live poliovirus exposure were fit by maximum likelihood assuming binomial sampling. Models for positive-definite quantities (concentration of poliovirus in stool, antibody titer) were estimated by ordinary least squares on log(quantity), and 95% confidence intervals assume log-normality. 95% confidence intervals were estimated by parametric bootstrap with 1000 replicates. To estimate bootstrap confidence intervals of parameters that are conditionally-dependent on previously estimated parameters, we propagated uncertainty by resampling known parameters from the 95% confidence intervals prior to resampling the data and re-estimating the parameters under investigation. Model equations and more detailed discussions of design decisions and calibration results can be found in S2 Text. Differences in comparable quantities are considered statistically significant at *α* = 0.05.

## Within-host model

Our within-host model describes shedding from and susceptibility to poliovirus infection. In the model, the ability of an infected individual to transmit polio depends on the duration of shedding and the concentration of poliovirus in their stool. Oral susceptibility to infection depends on the dose response relationship for the probability that poliovirus ingested orally results in an infection, as detected by subsequent fecal shedding. Shedding duration, concentration, and oral susceptibility all depend on pre-exposure immunity and the poliovirus source, vaccine or wild.

Immunity in our model is represented by the OPV-equivalent antibody titer (denoted *N*_Ab_)—an indirect measure of immunity that is inferred from measurements of shedding duration and/or dose response (first introduced by Behrend *et al* [25] and called “mucosal immunity” therein). Previous reviews have demonstrated that when immunity is due to prior OPV immunization or natural WPV infection, homotypic (of the same serotype) serum neutralizing antibody titers (measured as the geometric mean reciprocal dilution of serum that is able to neutralize 100 CID50 of poliovirus) are predictive of fecal shedding and susceptibility [25, 52]. However, serum antibody titers induced by IPV alone, and heterotypic titers against type 2 from bivalent type 1 and 3 OPV (bOPV), are not predictive of shedding and susceptibility [25, 53, 54]. The OPV-equivalent antibody titer describes the impacts of vaccination histories containing IPV or bOPV on shedding and susceptibility in terms of equivalent serum antibody titers from homotypic OPV vaccination in children. This model is agnostic about the biophysical mechanisms of immunity that prevent fecal shedding and is not intended to represent IgA concentration or other direct correlates of mucosal immunity [55]. Following the results of Behrend et al [25], we assumed that the typical immunologically-naive individual with no history of poliovirus exposure (“unvaccinated”) and no measurable humoral immunity (“seronegative”) has an OPV-equivalent antibody titer of *N*_Ab_ = 1 by definition, that the typical OPV-equivalent titer at maximum achievable individual immunity is *N*_Ab_ = 2048 (= 2^11^), and that homotypic antibody titers for each poliovirus serotype are independent.

The studies used to calibrate the within-host model span many countries, years, and types of immunization history. All included studies describe the fraction of subjects positive for poliovirus in stool after OPV challenge or WPV exposure as equal to the number of subjects shedding divided by the number tested at each time point. In many cases, the data were digitized from published figures that do not report variation in the number of samples for each time point, and so our sample sizes at each time point are often approximate. A summary of all included studies, with details about which studies contributed to which components of the model, and reasons for study exclusion appears in S2 Text [36, 41, 53, 54, 56–72], and the data are in S1 Structured Code and Data.

### Shedding duration

In Fig 1, we summarize available data for and our model of the impact of different immunization histories on shedding duration [36, 41, 53, 54, 56, 57, 59–62, 64–67, 69–71]. Fig 1A shows the average shedding duration survival distribution for all included trial arms by poliovirus strain (Sabin 1,2,3 or WPV) and pre-challenge immunization history (formulation and number of vaccine doses). All included subjects were five years of age or younger and from the Americas, Europe, the Middle East, or East Asia. We averaged across differences in vaccination schedule (i.e. tOPV at 6, 10, 14 weeks of age [54] was grouped with tOPV at 7, 8, 9 months [61]) and age at challenge because our goal was to model levels of immunity that describe typical conditions among children in practice—where vaccination schedules are not rigorously adhered to and natural oral challenge has no schedule. We also averaged over differences in the exact dose given (i.e. Salk vaccine [61] vs enhanced IPV [41]) as dose effects on shedding were insiginificant relative to differences in the type and number of vaccinations. At this stage in model building, we ignored waning and setting-specific variations in OPV take, both of which are addressed in later sections. Because individual-level data was not available, we could not construct proper Kaplan-Meier estimates of the survival distrubution for each immunization history. Rather, we assumed that the fraction shedding at each time point for each trial arm represented an approximate survival distribution, and the aggregated distributions shown in Fig 1 are the sample-size-weighted averages of the fraction shedding at each time point from the original papers. Thus, in rare instances where the sample sizes are small, the empirical distribution is not monotonically-decreasing as is necessary for a true survival model (i.e. Fig 1A: tOPVx2, Sabin 3). We used the data and our log-normal survival model for shedding duration (eq. (S1)) to inform estimates of typical OPV-equivalent antibody titers in children under five years of age with various immunization histories. Additional details about the data used, the shedding duration model, and model calibration can be found in S2 Text.

**Figure 1.**
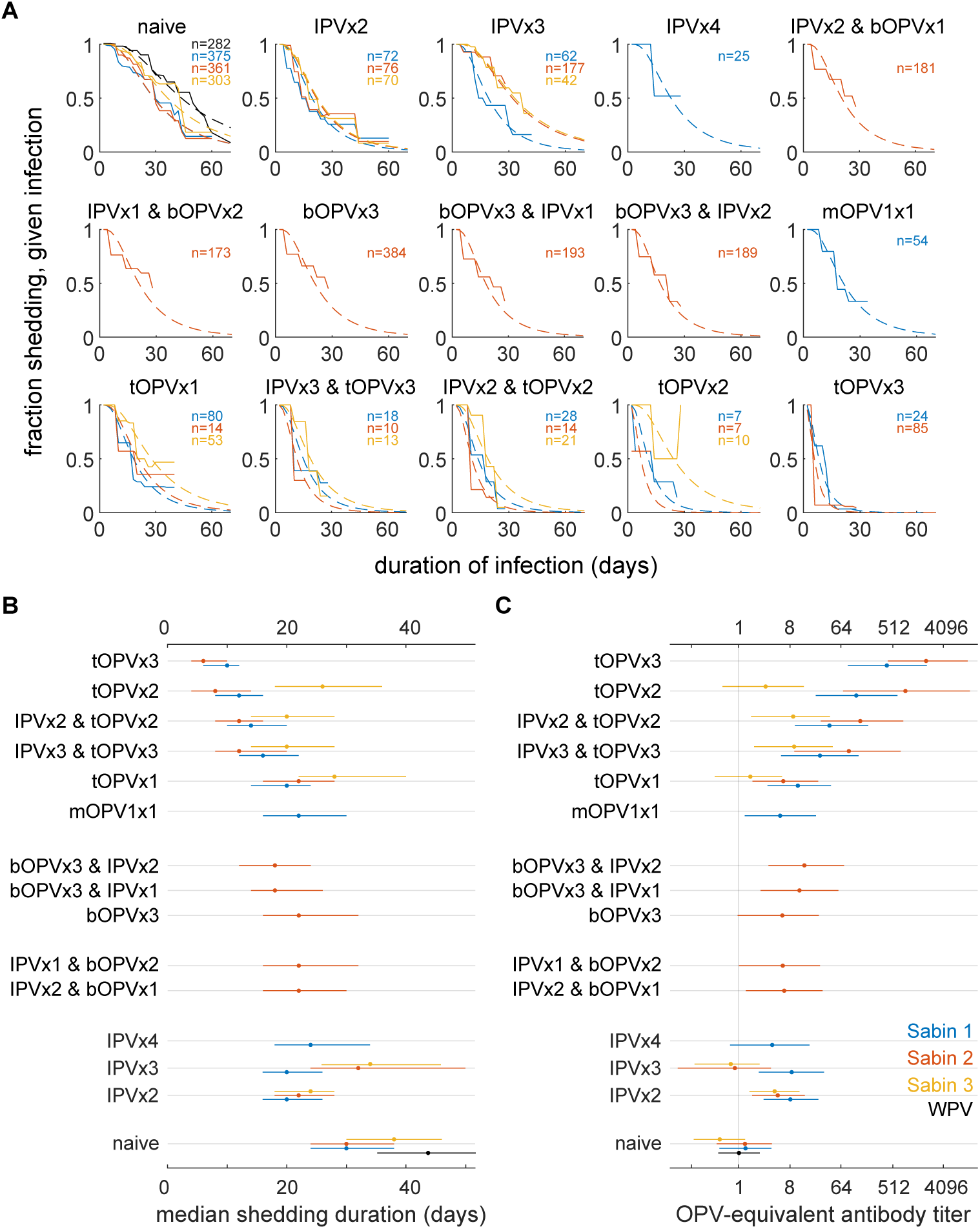
Effect of vaccination on shedding duration after OPV challenge and pre-challenge immunity. Labels describe vaccines received and number of doses (i.e. bOPVx2 & IPVx1: two doses of bOPV and one dose of IPV). (A) Shedding duration after OPV challenge: shedding duration survival curves from aggregated trial data (solid lines) and maximum likelihood model fit (dashed) for each immunization schedule and poliovirus type. (Infection with Sabin 1, blue; Sabin 2, red; Sabin 3, orange; WPV, black). (B) Median shedding durations after infection due to OPV challenge or WPV transmission estimated from model fits in panel A and (C) corresponding pre-challenge homotypic OPV-equivalent antibody titers. (See also S2 Text and an interactive visualization of shedding duration data is available at famulare.github.io/cessationStability/.)

Our shedding duration model (eq. (S1)) summarizes the following observations. In immunologically-naive individuals, there were no statistically significant differences by serotype in shedding duration after OPV challenge. The maximum likelihood estimate (MLE) of the median shedding duration for immunologically-naive individuals shedding any Sabin strain is 30.3 (23.6, 38.6) days, significantly shorter than the median shedding duration for WPV, 43.0 (35.7, 51.7) days (see also Fig S1). The median shedding duration associated with maximum OPV-equivalent antibody titer (*N*_Ab_ = 2048) is 6 (4, 10) days for the Sabin strains and is modeled to be 8 (6, 13) days for WPV. Repeated immunization with trivalent OPV (tOPV) has a cumulative effect on shedding duration. tOPV has weaker effects on shedding duration for type 3 than types 1 and 2, reflecting known per-dose efficacy differences [73]. At the level of aggregation examined here, limited data suggest mOPV is comparable to tOPV. Repeated vaccination with IPV alone shows no cumulative effect on shedding duration and with study-to-study variation showing little or no effect overall. bOPV produces a decrease in shedding duration against heterotypic Sabin 2 challenge, and this effect may be weakly enhanced by IPV after bOPV but not IPV before.

The transformation from median duration (Fig 1B) to titer (Fig 1C) serves as the definition of the OPV-equivalent antibody titer in our model—short post-challenge shedding duration implies high pre-challenge OPV-equivalent immunity. Having used the typical post-vaccination shedding duration distributions to define the OPV-equivalent antibody titer model, for the rest of this paper, we do not rely on immunization histories to determine immunity. Rather, we calibrate to setting-specific data on shedding duration to infer the appropriate OPV-equivalent immunity for each data source. This allows us to generalize from the aggregated results in Fig 1 to incorporate variation in post-vaccination immunity without having to model mechanisms of variation, such as enterovirus interference or enteropathy [25, 74–77].

### Concentration of poliovirus in stool

In Fig 2, we show available data for and our model of the concentration of poliovirus in stool while shedding after OPV challenge [53, 54, 56–58, 67, 68]. The included studies all reported concentration as the geometric mean 50% culture infectious dose per gram of stool (CID50/g) averaged across all subjects positive for poliovirus at each time point, and individual-level variation was generally not reported. Ages at OPV challenge ranged from 6 months to 65 years or more. The majority of trial arms challenged subjects with mOPV2 (mOPV1, *n* = 5; mOPV2, *n* = 11; mOPV3, *n* = 5). Data exploration revealed no systematic differences in concentration by serotype. We are not aware of similar longitudinal data for WPV shedding. OPV-equivalent antibody titers were estimated from the corresponding shedding duration data for each trial arm (see S2 Text), and trial arms considered immunologically-naive reported no history of live poliovirus exposure, contained confirmed seronegative subjects, or had OPV-equivalent antibody titers consistent with *N*_Ab_ = 1.

**Figure 2.**
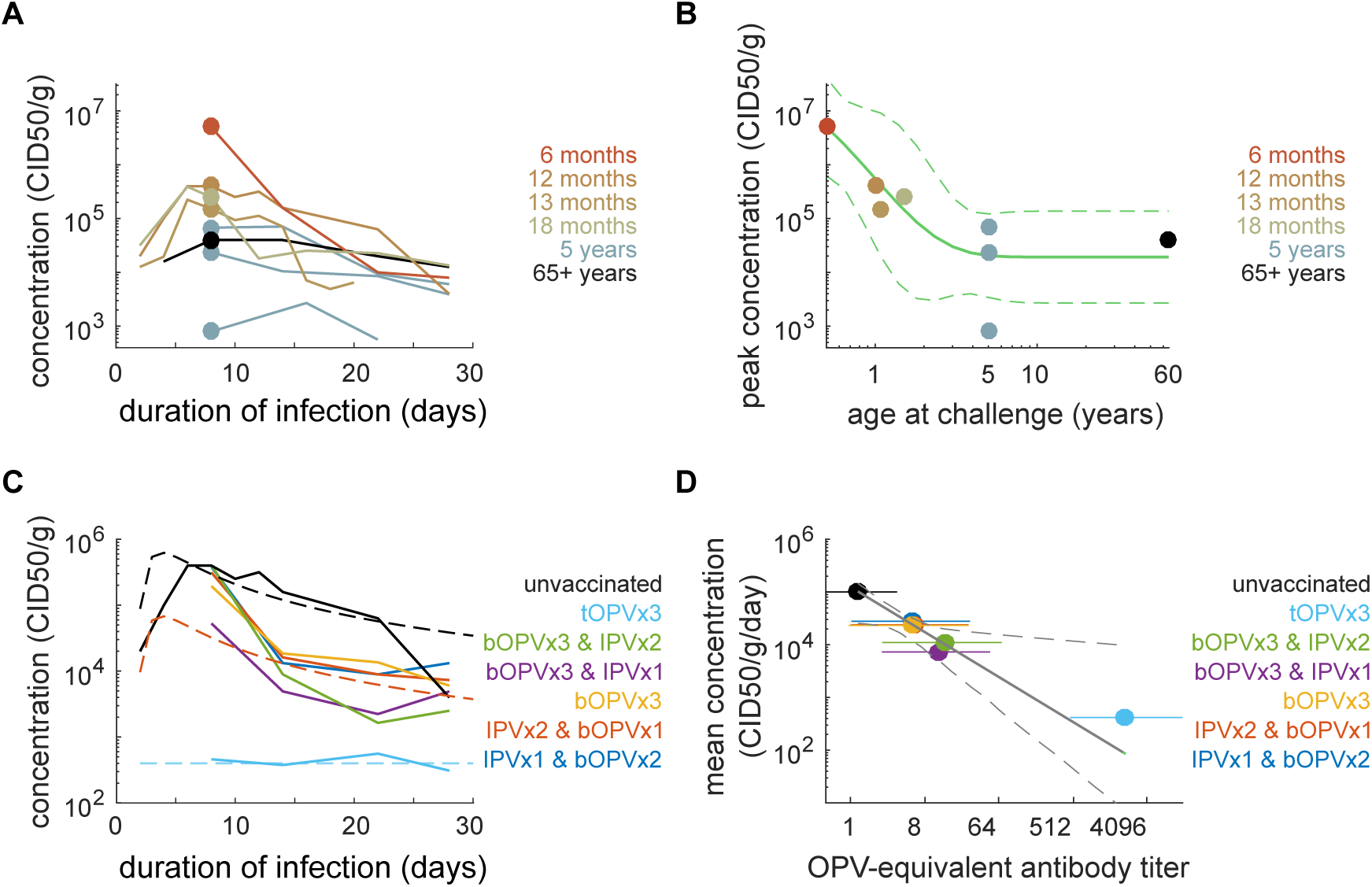
Concentration of poliovirus in stool: effects of age and immunity. (A) Mean concentration of polivirus in stool (CID50/g) vs. time after OPV challenge for immunologically-naive subjects (color by age at challenge). (B) Peak concentration depends on age (dot color by age at challenge, corresponding to data from panel A at one week post-challenge; green line, model MLE and 95% CI, eq. (S2)). (C) Mean concentration after mOPV2 challenge for subjects with various vaccination histories (dashed, model MLE; solid, trial data age-adjusted to 12 months using eq. (S3)). The concentration of poliovirus in stool depends on pre-challenge vaccination history. (D) The mean daily concentration (culture infectious doses per gram per day; CID50/g/day) declines by one order of magnitude for every eight-fold increase in OPV-equivalent antibody titer (OPV-equivalent titers (color, MLE and 95% CI) for each trial arm shown in panel C; black, model (eq. (S4), MLE and 95% CI)). Interactive data visualization available at famulare.github.io/cessationStability/.

Our model of poliovirus concentration in stool summarizes the following observations. Poliovirus concentrations peak 5–8 days after acquiring infection and decline slowly thereafter (Fig 2A). Data from immunologically-naive subjects revealed an unexpected dependence of peak concentration with age (Fig 2B). Peak concetration declines by roughly two orders of magnitude over the first three years of life with an exponential aging constant of 12 (1, 45) months, consistent with major developmental milestones including the transition to solid food and immune system maturation [78, 79], after which the limited data indicate stability of peak shedding concentration for life. The concentration of poliovirus in stool is correlated with vaccination history (Fig 2C) and decreases by roughly a factor of ten with each eight-fold increase in OPV-equivalent titer (Fig 2D). The shedding concentration model is described in eqs. (S2–S4).

### Oral susceptibility to infection

To inform our dose response model for oral susceptibility to infection, we first examined studies of healthy children that measured the probability of fecal shedding after receiving oral droplets with doses ranging from 10^1^ to 10^6^ CID50, and for which pre-challenge immunization histories were known. Three studies challenged with Sabin 1 [41, 61, 63], none used Sabin 3 or WPV, and one study challenged with Sabin 2 and type 2 poliovirus derived from Sabin 2 after five days of replication in vaccinated children [58]. There were no statistically significant differences between Sabin 1 and Sabin 2 across these trials, but statistical power at low doses is poor. We also included modern studies of vaccine doses (10^5−6^ CID50) that provided information about the effects of heterotypic immunity against type 2 from bOPV [53, 54] and IPV boosting on prior OPV immunization [72]. OPV-equivalent antibody titers were estimated from the corresponding shedding duration data for each trial arm.

Our dose response model summarizes the following observations (Fig 3A-B). The typical Sabin 1 dose required to infect 50% of immunologically-naive healthy children (the 50% human infectious dose, HID50) is 54 (26, 100) CID50, and the fraction shedding approaches one for doses greater than 104 CID50. Immunity has similar effects on susceptibility as it does on shedding duration and concentration. IPV-only immunization reduces susceptibility to infection in some studies but not all, and the effect is at most comparable to that provided by heterotypic immunity against type 2 from immunization with bOPV. tOPV reduces susceptibility across all doses. Not addressed in previous sections on shedding is that IPV-boosting in subjects with prior OPV immunization is highly effective for reducing susceptibility—as is now well-known [72, 80, 81]. OPV-equivalent antibody titer has a monotonic relationship with oral susceptibility (Fig 3C). The data are consistent with an immunity-dependent beta-Poisson dose response model [82] (Fig 3D and eq. (S5)).

**Figure 3.**
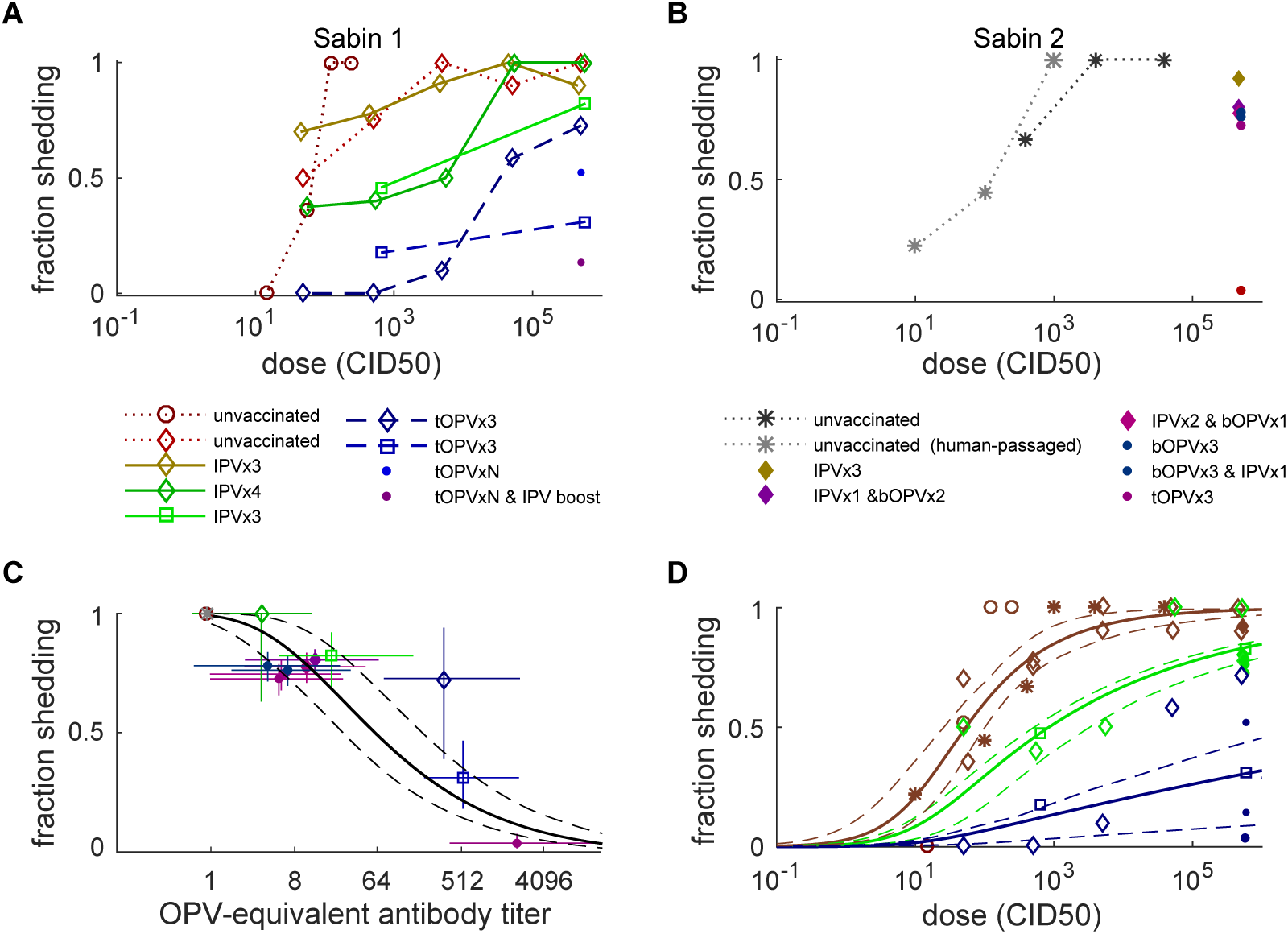
Oral susceptibility to infection after OPV challenge. (A) Fraction shedding after Sabin 1 oral challenge at different doses (color by trial arm, symbol by source study). (B) Fraction shedding after Sabin 2 oral challenge at different doses (color by trial arm, symbol by source study; data for doses ≤ 10^3^ culture infectious doses (CID50) are for human-passaged Sabin 2 isolated from stool five days after vaccination). (C) Fraction shedding at vaccine doses (10^5−6^ CID50) decreases with increasing OPV-equivalent antibody titer. (Color and symbols as in panels A–B; black lines are model MLE and 95% CI using eq. (S5)). (D) Beta-Poisson dose response model MLE and 95% CI. Three model scenarios shown correspond to immunologically-naive (*N*_Ab_ = 1, red), heterotypic bOPV and upper-bound IPV-only (*N*_Ab_ = 8, green), and typical tOPV or post-IPV-boosting (*N*_Ab_ = 256, blue). Data from panels A–B (symbols as above, colored by corresponding model scenario).

To inform our model of strain-specific differences in dose response, we examined two transmission studies from similar settings in the United States. The first study in Houston in 1960 [36] measured transmission among immunologically-naive close contacts of vaccinees for each of the Sabin strains, and another in Louisiana from 1953–55 [35] measured close contact transmission of WPV (combined across all serotypes); these studies and our transmission model are described in detail in a later Methods section. Under the assumptions that Sabin 1 is well-described by the OPV challenge model above and that sanitation and contact patterns are similar across the four trial arms, differences in transmission are attributable to the virus-specific differences in infectivity shown in Table 1.

**Table 1.** Infectivity by strain. MLE and 95% CI of the HID50—the oral dose that infects 50% of immunologically-naive people.

| strain | HID50 |
| --- | --- |
| Sabin 1 | 54 (26, 100) TCID50 |
| Sabin 2 | 30 (15, 54) TCID50 |
| Sabin 3 | 67 (34, 120) TCID50 |
| WPV | 7 (2, 41) TCID50 |

### Waning immunity

We built a composite picture of waning immunity against infection from analysis of OPV-equivalent antibody titers across studies. We considered data for individuals that were likely maximally immune after their last poliovirus exposure, either due to immunization with three or more doses of tOPV [41, 54, 72] or accumulated natural immunity through 15 years of age during the endemic era [56, 68]. The included trial arms involved subjects from 6 months to 65+ years of age and with between 1 month and likely 45+ years from last immunizing event to OPV challenge (see S2 Text for additional details).

Our waning model summarizes the following observations (Fig 4). Absent reinfection or vaccination, immunity declines over many years, possibly with increasing variation in adults. We modeled waning as a power-law decay [83] during the months since last immunization, *N*_Ab_(*t*) ∝ *t*^−*λ*^, with exponent *λ* = 0.87 (0.73, 1.02) (eq. (S6)). The limited relevant data after bOPV vaccination [54] are consistent with the hypothesis that heterotypic and homotypic immunity share waning dynamics (bOPV data only, *λ* = 0.52 (0, 1.2)).

**Figure 4.**
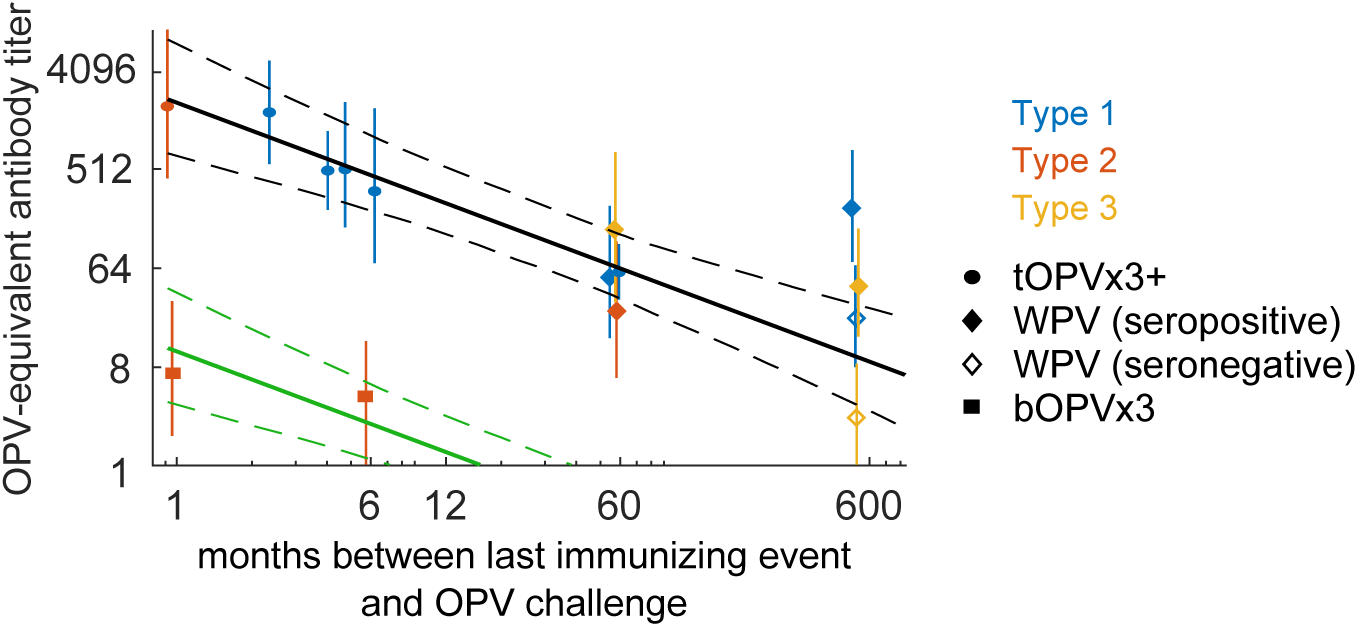
Waning immunity against infection. OPV-equivalent antibody titer vs. time between last exposure and mOPV challenge (color by serotype and symbol by source of immunity). Power-law model of waning from peak homotypic immunity (MLE and 95% CI, black lines) and heterotypic immunity against type 2 from bOPV (MLE and 95% CI assuming homotypic waning exponent, green lines).

### Transmission model

Our model describes the effects of within-host dynamics on transmission among people who share a household and close social contacts outside the household. We assumed that transmission from infected person to recipient occurs by oral exposure to infected feces, where the amount of poliovirus transmitted per exposure is determined by the shedding duration and concentration models, and recipient susceptibility is determined by the dose response model.

### Person-to-person transmission

The population structure of the model is based on the essential transmission network motif examined by the field transmission studies (Fig 5A): an index person transmits to household contacts (typically family members), who in turn transmit to their close social contacts outside the household. For each of the three individuals along the transmission chain, the person-to-person model calculates daily incidence (the probability of becoming infected each day), prevalence (the probability of shedding poliovirus in stool each day), and concentration of poliovirus shed (CID50 per gram of stool).

**Figure 5.**
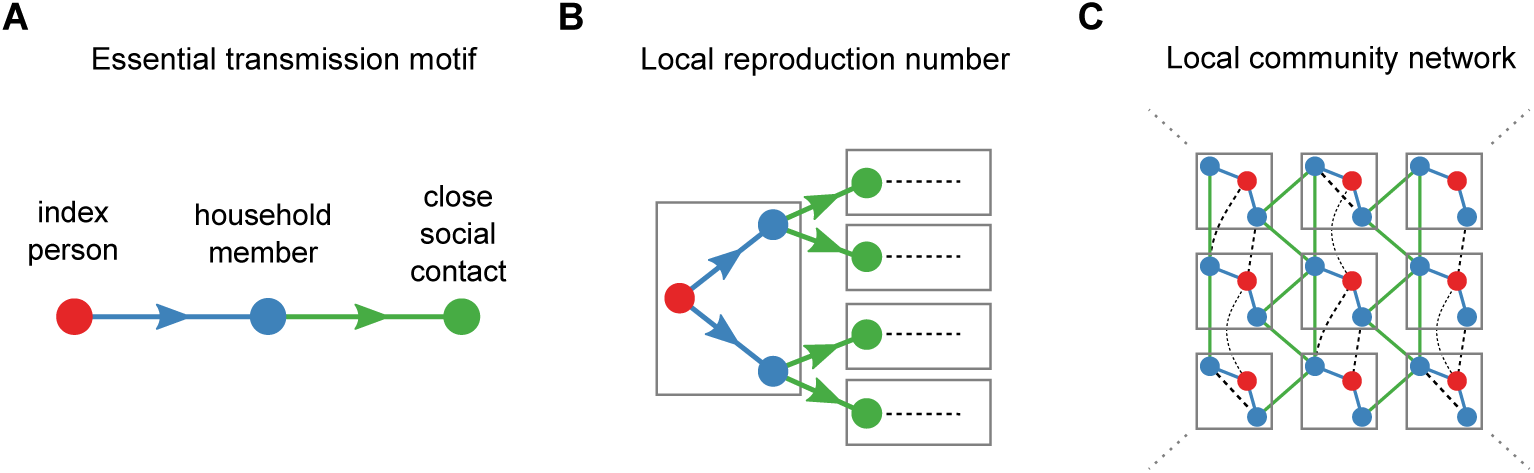
Network motifs of poliovirus transmission. (A) The essential motif of poliovirus transmission is index person to household member to close social contacts (dot color gives subject type, line color describes relationship). (B) The local reproduction number describes the expected number of secondary households infected by an index person based on sanitation, individual immunity, and the number of close social contacts. The house-to-house transmission motif represents this first generation of local transmission (color as in panel A; gray boxes denote households; dashed lines indicate relationships beyond the first generation). (C) Our definition of the local reproduction number captures transmission among household members and close social contacts (solid colored lines) but does not include all relationships that may contribute to transmission (dashed black).

Our model focuses on person-to-person transmission because it allows us to study factors that causally affect transmission probability within and between households with contact tracing data collected for that purpose (as described in detail below). Although we rely in this paper on calibration to poliovirus transmission data, each parameter has biophysical meaning and can in principle be measured directly in the absence of live poliovirus. This model building approach offers a complementary alternative to more classical models that focus on population-wide measures of disease transmission and for which key transmission rate parameters lack biophysical meaning [84]. Our focus on specific within and between household relationships follows the available data and emphasizes the roles of the strongest links in the transmission network to determine community susceptibility to poliovirus transmission.

Infections in index persons are defined to begin on day *t* = 1 due to either mOPV or WPV exposure on day *t* = 0. Incidence is determined by the dose response model,

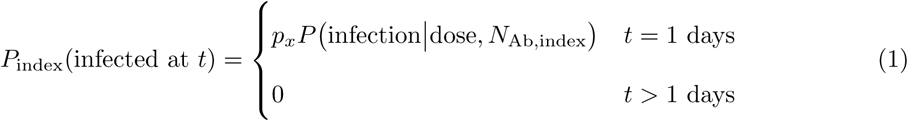

where *p*_*x*_ (serotype *x* = 1, 2, 3, *p*_*x*_ ∈ (0, 1]) is a setting-specific susceptibility modifier that accounts for non-immunological host factors such as non-polio enterovirus infection or enteropathy that can reduce the probability of shedding [25, 74, 77], and the second term is defined in eq. (S5). The prevalence for *t* > 0 after exposure is given by:

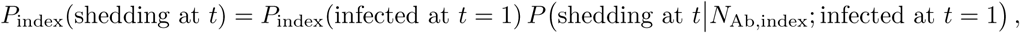

where the first term is eq. (1) and the second is the shedding duration model in eq. (S1). Household members are infected with probabilities determined by the dose response model, the size of the fecal dose, and the amount of virus shed by the index person. Daily incidence derives from exposure to index shedding as:

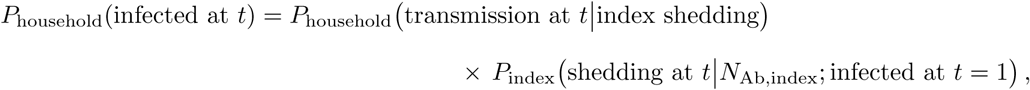

with

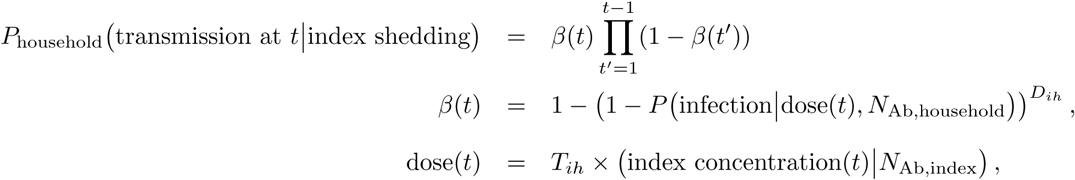

where *P*_household_ transmission at *t* index shedding is the household member incidence on day *t* given contact with a shedding index person, *β*(*t*) is the infection probability determined by the dose response model, *D*_*ih*_ is the interaction rate for an index and household member pair (average number of fecal-oral exposures per day), *T*_*ih*_ is the fecal-oral dose (micrograms of stool per exposure), and index concentration (CID50 per gram) is given by the fecal concentration model in eq. (S4). Household member prevalence follows from convolving daily incidence (assuming no re-infection) with the shedding duration distribution:

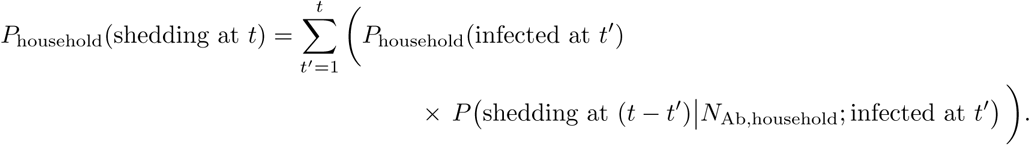

The model assumes all transmission to close social contacts occurs only through household members of index cases, depending on contact susceptibility and fecal exposure to and the amount shed by the contacted household member. Daily incidence derives from exposure to household contact shedding as:

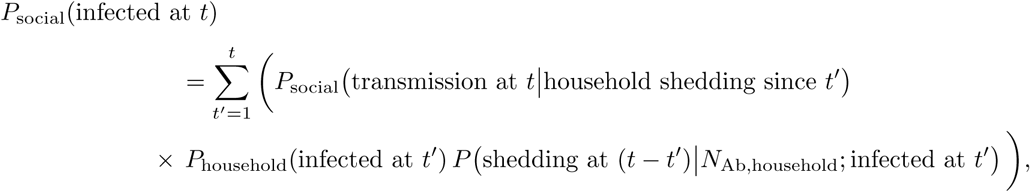

with

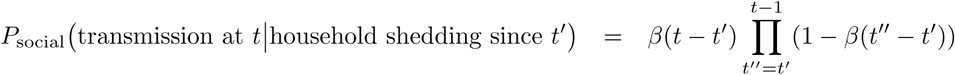

and

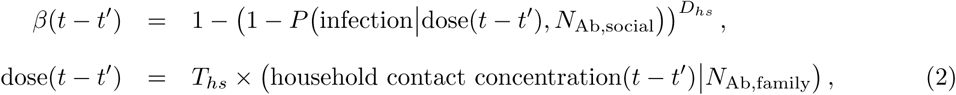

where *D*_*hs*_ is the interaction rate for a household-to-close social contact pair, *T*_*hs*_ is the fecal-oral dose, and (*t* − *t′*) is the time interval since the household contact became infected. The convolution over household member incidence accounts for all the times at which household members can become infected. Close social contact prevalence follows from convolving daily incidence with the shedding duration distribution:

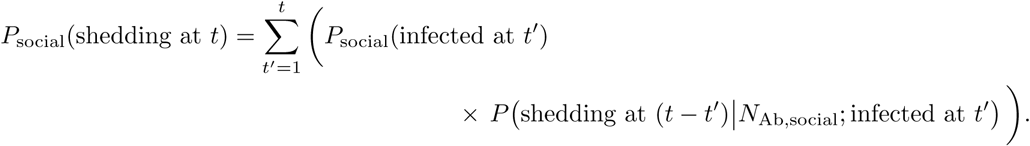

### Local reproduction number

We defined the local reproduction number (*R*_loc_) as the expected number of close social contacts infected by an index person due to transmission along the index-household-social contact essential transmission motif,

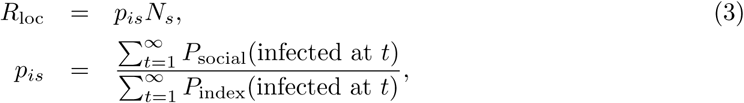

where *N*_*s*_ is the average number of close social contacts outside the index household and *p*_*is*_ is the total probability that an index person transmits through a household member to a close social contact in another household, as determined by the ratio of total incidences in eqs. (1) and (2). *R*_loc_ describes the first generation of household-to-close social contact transmission following infection of an index person (Fig 5B).

Within close-knit communities where all households have similar demographic, behavioral, and immunological patterns, *R*_loc_ provides a lower bound on the total transmission rate because it does not include all possible transmission routes (Fig 5C). Across large, heterogeneous communities, the *a priori* relationship between *R*_loc_ and the true average transmission rate across all contacts is unclear. The model can be extended to describe any set of relationships—for example, a household member may have many more socially-distant contacts that receive smaller doses less often—but the model complexity that needs to be constrained increases rapidly with the number of relationships. For these reasons, this iteration of the model cannot make predictions about the absolute probability or severity of outbreaks, for which model specification is critical [46–50]. Rather, *R*_loc_ is a useful threshold parameter for categorizing outbreak risk with data from contact-tracing studies.

### Calibration

While the data on within-host aspects of polio infection showed remarkable coherence across studies from different eras and settings, this is not the case for literature on community transmission of poliovirus. The eighteen transmission studies reviewed by Tebbens *et al* [24] exhibit varying thoroughness in their reporting pre-exposure immunity and contact relationships. In lieu of a comprehensive review, we based our transmission model on specific studies capable of identifying important model parameters. The studies took place in the United States between 1953 and 1960 [35, 36] and India between 2003 and 2008 [37]; all had large sample sizes, carefully reported demographic and social contact attributes, and provided sufficient information to infer pre-exposure OPV-equivalent immunity (either directly through vaccination histories or serostatus, or indirectly via shedding duration). The fraction of subjects positive for poliovirus after OPV challenge or WPV exposure was given by the number of subjects shedding in stool [36, 37] or recently seroconverted [35] divided by the number tested. Additional information about calibration methods are provided in S2 Text.

We assumed that the serotype-specific dose response model parameters (Table 1, eq. (S5)) are independent of setting. The setting-specific free parameters are the pre-challenge OPV-equivalent antibody titers for each subject type (*N*_Ab_), the average fecal-oral dose (micrograms of stool ingested per interaction, *T*_*ih*_ and *T*_*hs*_), the interaction rates (number of fecal-oral contacts per day for each person-to-person pair, *D*_*ih*_ and *D*_*hs*_), the setting-specific dose response modifiers (*p*_*x*_), and the typical number of close social contacts (*N*_*s*_). The interaction rate and fecal-oral dose parameters are not separately identifiable from the available data, and so we fixed the index-to-household-member interaction rate to once per day (*D*_*ih*_ = 1) and assumed that fecal-oral dose is independent of relationship type (*T*_*ih*_ = *T*_*hs*_).

From the Sabin transmission study conducted in Houston 1960 [36], we calibrated the serotype-specific dose response parameters, and the fecal-oral dose and between-household interaction rate representative of a typical endemic setting with low socioeconomic status in the pre-elimination United States. Additional study-specific parameters described OPV-equivalent immunity and trial-to-trial variation in post-vaccination shedding in index children. Briefly, children aged 2 to 18 months were enrolled to receive a dose of mOPV. Weekly stool samples were collected from the vaccine-recipient index children, their siblings, and primary extrafamilial social contacts of siblings. The majority of index children had prior serological immunity either due to maternal antibodies or IPV vaccination. Pre-challenge serology was not presented for siblings or extrafamilial contacts. The authors observed no significant differences in shedding by IPV immunization history or pre-challenge serologic immunity. Family members and extrafamilial contacts five to nine years of age shed significantly less from transmission, and there was essentially no shedding in subjects older than ten years of age (S2 Text). From joint calibration across the three mOPV trial arms (Fig 6A), we inferred that children under five years of age who shed poliovirus, regardless of position in the transmission chain, had OPV-equivalent antibody titers of *N*_Ab_ = 1, and the fraction of infants shedding one week after receiving mOPV was high: type 2, 0.92 (0.85, 1.0), type 1, 0.79 (0.70, 0.88), and type 3, 0.81 (0.71, 0.91). Thus it is likely that most children under five had no experience with WPV. (See S2 Text for additional details.) From the differences in transmission by serotype in this immunologically-naive population, we estimated the infectiousness of each serotype (shown above in Table 1). The estimated fecal-oral dose was microscopic at 5 (1, 31) micrograms per day (*µ*g/day), and the estimated interaction rate in a family member and extrafamilial contact pair—the average number of fecal-oral exposures per day—is 9.0 (2.6, 46), possibly reflecting higher rates of social interaction in peer versus infant-sibling pairs [85].

**Figure 6.**
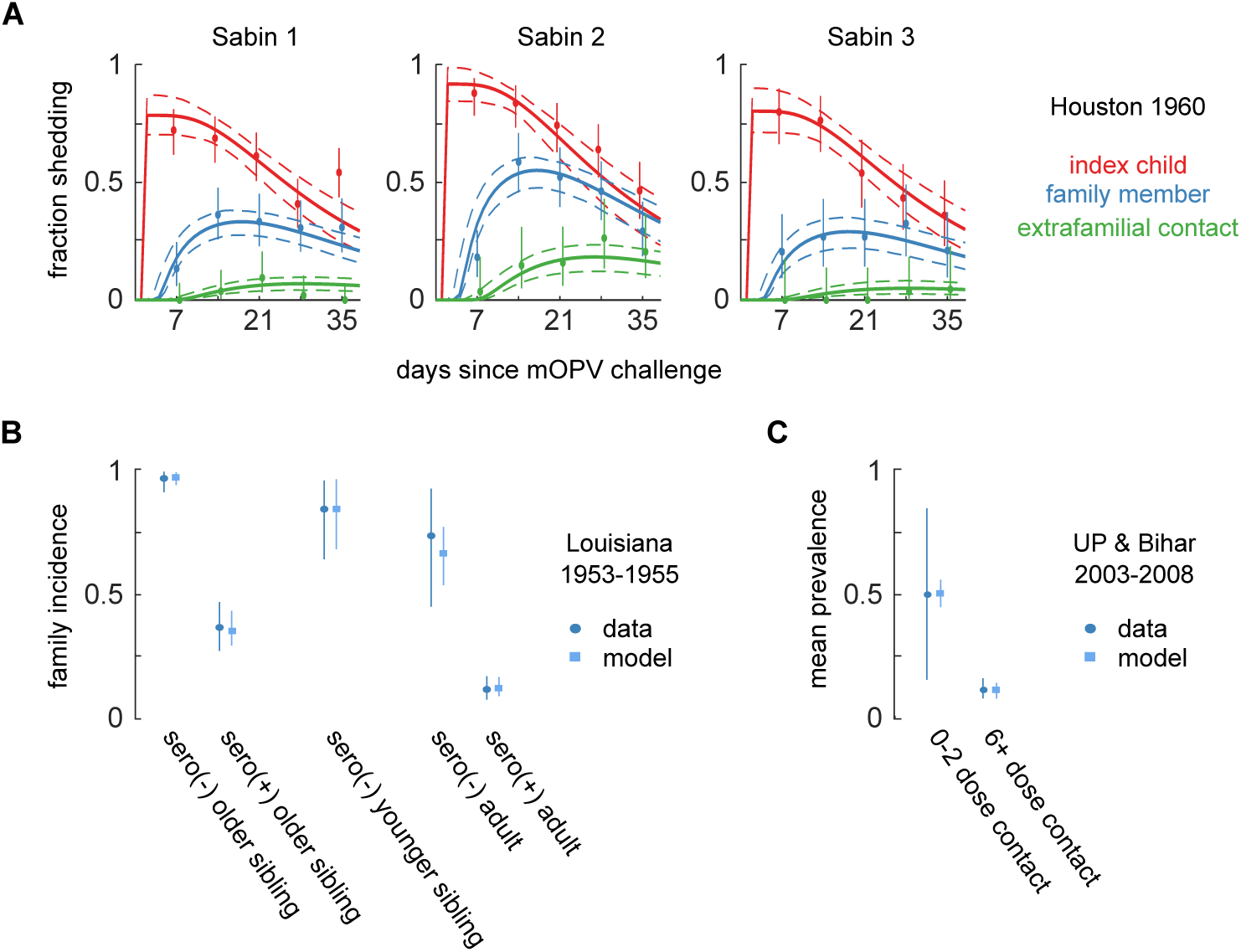
Transmission model calibration. Each study measured the amount of transmission from index persons in different ways. (A) Houston 1960: fraction of children under five years of age shedding each week after mOPV challenge to an index child and subsequent transmission. (Color by subject type; weekly data MLE and 95% CI, dot-and-whiskers; model MLE and 95% CI, lines). Eight free parameters are jointly identified across the nine calibration targets. (B) Louisiana 1953–1955: incidence in household contacts of index children naturally infected by WPV, measured by seroconversion approximately 30 days after the index child became infected. Three free parameters are jointly identified by the five calibration targets. (C) Uttar Pradesh & Bihar 2003–2008: mean prevalence of WPV in stool measured in household contacts after the onset of paralysis in index children. One free parameter is jointly identified by the two calibration targets. See S2 Text for additional information about model fit.

From the WPV transmission study conducted in Louisiana from 1953–1955 [35], we calibrated WPV dose response and the age-dependence of the fecal-oral dose, under the assumption that the fecal-oral dose between index children and older siblings was the same as in Houston 1960. Briefly, Gelfand *et al* enrolled families with newborn children to undergo monthly surveillance for naturally-acquired polio infections. Whenever a newly-infected index child was identified, household contacts were tested for subsequent polio infection, most reliably through evidence of seroconversion. This measure of incidence was reported for siblings and parents, stratified by serostatus and age relative to the index child. We assumed that the OPV-equivalent antibody titer of seronegative subjects was *N*_Ab_ = 1, and we reconstructed from the published serological data that the median seropositive titer was *N*_Ab_ = 93 in this naturally-immunized population. Joint calibration of incidence thirty days after index infection across the five reported index-family relationships (Fig 6B) confirmed the expected outcome that WPV is more infectious than any Sabin strain (Table 1). We inferred that the fecal-oral dose transmitted from index children to adults was 26 (16, 41)% of that passed to siblings under age five; a similar age-related decline in fecal-oral dose was inferred with this same model for a recent Sabin 2 transmission study in Bangladesh [86]. The estimated fecal-oral dose transmitted from older index children to younger siblings was 46 (26, 104)% of the reverse.

To estimate an upper-bound for fecal-oral dose in regions of extremely high polio transmission intensity [87], we examined WPV surveillance data from 2003–2008 in India and reported by Grassly et al [37]. The authors examined the fraction of stools positive for WPV from children under five years of age who were household contacts (siblings, residents of the same household, or playmates [37]) of paralytic WPV cases (mostly from Uttar Pradesh (UP) and Bihar). Household contacts with low immunity (0–2 reported tOPV doses) and high immunity (6+ reported doses) were grouped for analysis. They estimated that 51% (16, 84)% of low immunity and 12 (8, 16)% of high immunity contact stool samples were positive for WPV when sampled once during the ten weeks after paralysis of the index child. For our model, we assumed that the high immunity cohort had an OPV-equivant antibody titer of *N*_Ab_ = 512, corresponding to their estimate of an eleven day mean shedding duration, and that the low immunity cohort had *N*_Ab_ = 1 in this setting known for low tOPV efficacy [75]. Given the assumptions, and after accounting for the unobserved time infected prior to paralysis (see S2 Text), we inferred from joint calibration to both targets (Fig 6C) that the fecal-oral dose transmitted from index children to household contacts 230 (2, 1800) *µ*g/day, roughly fifty times higher than in Houston 1960.

### Additional assumptions

The calibration studies did not report sufficient information to constrain the average number of close social contacts outside the household (*N*_*s*_), and only the Houston study provided information about the household-to-close social contact interaction rate (*D*_*hs*_). Except when explicitly exploring sensitivity to these parameters, we made the following assumptions. For Houston/Louisiana, we assumed that the typical number of close social contacts is *N*_*s*_ = 4 (3, 5), reflecting the average number of close friends in American childhood social networks [88]. For UP and Bihar, we assumed *N*_*s*_ = 10 (8, 12), based on scaling Houston in proportion to the two-to-three times larger typical classroom sizes [89, 90] and population densities [91, 92] in northern India. For all settings, the value for the household-to close social contact pair interaction rate (*D*_*hs*_) estimated from Houston was used.

To simplify the presentation of results below, we chose to ignore adults. First, calibration showed that changes in childhood immunity from vaccination policy choices have larger effects on immunity than waning (Fig 1C and Fig 4), and so typical adults alive near OPV cessation will make small contributions to the local transmission rate relative to children. Second, unimmunized adult family members of infected children have similar (albeit slightly lower) likelihood of infection from index persons than unimmunized children (Fig 6B) [35, 86]. Absent immunity, including adults in our model is roughly equivalent to increasing the number of child contacts.

## Results

Fig 7 summarizes our within-host model for the effects of immunity on shedding and susceptibility and how typical immunity levels relate to specific vaccination schedules. The shedding index (Fig 7A) is the expected total amount of virus shed per gram of stool after mOPV challenge. For a typical healthy child under five years of age—averaged over vaccination timing and waning—each of the first three doses of OPV increases the OPV-equivalent antibody titer by roughly a factor of eight and decreases the expected amount of virus shed by a factor of ten. To characterize settings with low OPV effectiveness [25, 74–77], we found that children who received at least six doses of tOPV in Uttar Pradesh and Bihar [37] had similar OPV-equivalent antibody titers to healthy clinical trial subjucts who received three tOPV doses. In our model, IPV boosting and OPV doses after the first three maintain maximum immunity. The heterotypic protection against type 2 from bOPV immunization is comparable to that of a single homotypic dose but does not accumulate with multiple doses. We inferred from the trial arms reviewed that the OPV-equivalent immunity of IPV-only is at most comparable to heterotypic immunity from bOPV, but we expect that the true impact is closer to none—the trial arms that showed the highest immunity (Fig 1) likely included some incidental IPV boosting, with larger effects in older [41, 61, 65, 69] vs. younger [53, 57, 61] subjects in OPV-using countries, and negligible effects in older subjects in countries where OPV is not ubiquitous [67, 93]. Susceptibility is also strongly impacted by immunity, with the expected fraction shedding after Sabin 2 challenge dropping below half at all relevant doses for *N*_Ab_ *≥* 64 (Fig 7B).

**Figure 7.**
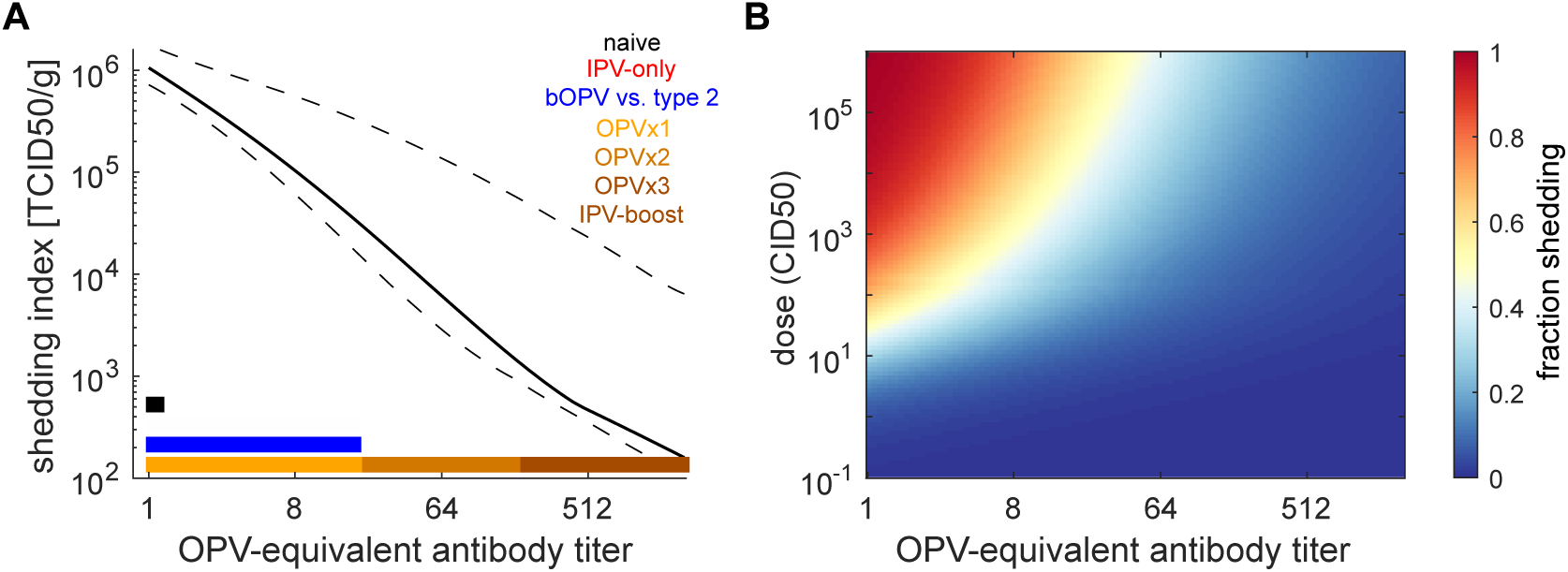
The effects of pre-exposure immunity on shedding and oral susceptibility to infection in children. (A) Shedding index vs. pre-challenge OPV-equivalent antibody titer. (Black, model MLE and 95%CI; color, range of immunities expected from vaccination.) In the legend, the number of OPV doses equivalent to a titer range assumes healthy child (typical clinical trial) vaccine take rates. (B) Dose response model vs. OPV-equivalent titer (dose measured in culture infectious doses (CID50)). Pre-exposure immunity reduces susceptibility at all doses.

Our waning model (eq. (S6), Fig 4) predicts that without reinfection, typical peak OPV-equivalent antibody titers (*N*_Ab_ = 2048) decline to typical three-dose healthy child immunity (*N*_Ab_ = 512) in 5 (4, 7) months and to typical two-dose immunity (*N*_Ab_ = 64) in an additional 4 (2, 10) years. However, the model also predicts that it takes an additional 45 (15, 160) years to fall to the equivalent of one-dose childhood immunity (*N*_Ab_ = 8) and that residual immunity persists for life, as has been suspected previously [24, 94]. This result disagrees with the conclusions of Abbink et al [68]. They argued from the lack of correlation between serological boosting responses and shedding duration after OPV challenge that memory immunity in seronegative elderly does not protect against poliovirus shedding, but the study lacked a control group of never-exposed subjects to contrast deeply waned and truly naive immunity. As seen through metastudy, the OPV-equivalent immunity of the Abbink *et al* seronegative elderly cohorts is similar to that of children who have received one dose of OPV. For heterotypic immunity against type 2 from bOPV, we predict that protection from shedding will be lost 13 (9, 22) months after bOPV vaccination is stopped [95].

Fig 8 shows maximum likelihood estimates from our transmission model for the local reproduction number of WPV, *R*_loc_ (eq. 3), as functions of immunity and daily fecal-oral dose (Fig 8A), and fecal-oral dose and the number of close social contacts outside the household (Fig 8B). The value of *R*_loc_, a measure of the average transmission rate in a community, depends linearly on the number of social contacts, but varies across four orders of magnitude due to strong effects of immunity and dose. Assuming one fecal-oral exposure per day (see Methods: Transmission model: Calibration), the physiological range for the average fecal-oral dose maxes out at two milligrams of stool, corresponding to the upper bound of our estimate from Uttar Pradesh and Bihar in 2003–2008. When all children have typical three-dose childhood immunity or more (*N*_Ab_ *≥* 512), we estimated *R*_loc_ *<* 1 over the entire physiological range, and thus that WPV persistence is impossible under universal tOPV immunization. In the absence of immunity, WPV epidemics are possible in all settings where sanitation practices permit the ingestion of roughly one microgram of stool per day or more.

**Figure 8.**
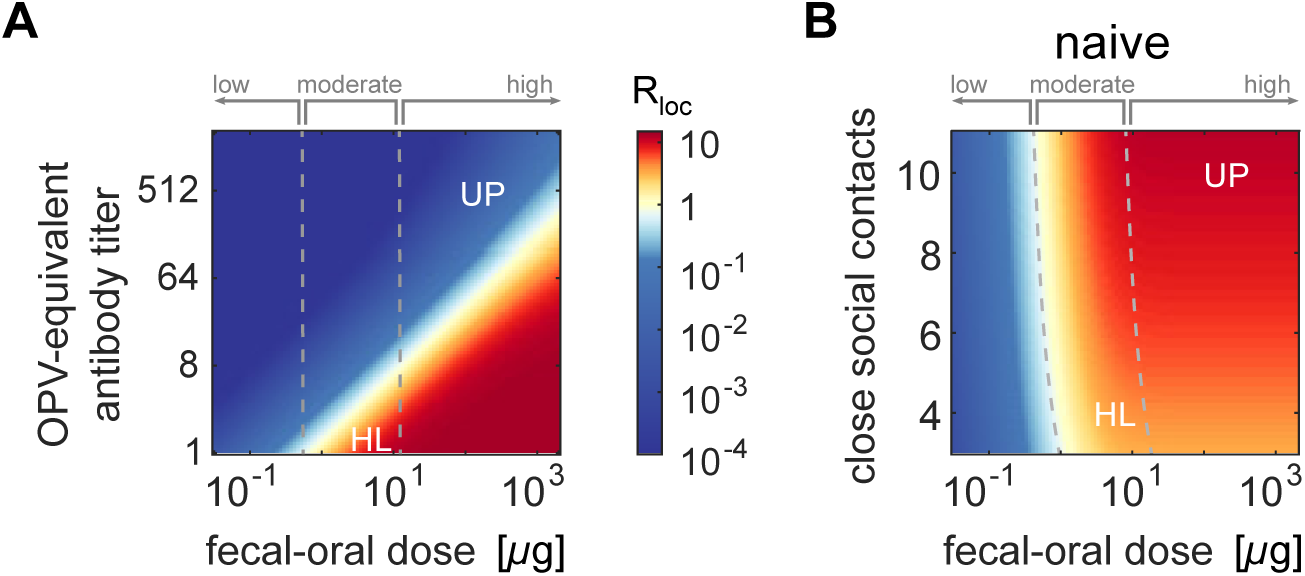
WPV local reproduction number depends on immunity, sanitation, and contact network size. (A) Local reproduction number vs. immunity and fecal-oral dose (assuming twelve close social contacts outside the household per index child and that everyone has equal immunity). (Colormap, *R*_loc_; dashed lines, transmission rate category boundaries; HL, Houston/Louisiana; UP, Uttar Pradesh and Bihar.) (B) Local reproduction number vs. fecal-oral dose and number of close social contacts (assuming all are immunologically-naive; legend as in panel A).

We identified three categories describing the transmission rate in different settings: low, where the fecal-oral route alone cannot sustain WPV transmission (*R*_loc_ *<* 1 for all *N*_Ab_ *≥* 1); moderate, where WPV epidemics can occur in immunologically-naive communities but not where at least one-dose OPV-equivalent immunity is common (*R*_loc_ *≥* 1 only for *N*_Ab_ *<* 8); and high, where WPV can persist despite at least one-dose OPV-equivalent immunity in everyone (*R*_loc_ *≥* 1 when *N*_Ab_ *≥* 8 but less than a protective threshold).

Fig 9 shows the dependence of the local reproduction number on poliovirus strain and immunity for example low, moderate, and high transmission rate settings. In low transmission rate settings, epidemic transmission of any strain cannot occur without contributions from the unmodeled oral-oral transmission route. This result supports the long-held hypothesis that oral-oral transmission is critical in settings with good sanitation, supported by many observations that IPV alone—an effective intervention against oral shedding [38–42]—can block transmission and prevent outbreaks from importation in communities with high socioeconomic status [8, 42, 96]. In moderate transmission rate settings (such as Houston 1960 [36], Louisiana 1953–1955 [35], or Matlab, Bangladesh 2015 [86]), immunologically-naive populations can support WPV epidemics, but *R*_loc_ ≲ 1 for the Sabin strains, and one-dose OPV-equivalent immunity (*N*_Ab_ = 8) is sufficient to block epidemic transmission of all strains. This result is consistent with the historical experience in middle- and high-development countries that WPV elimination rapidly follows the introduction of OPV vaccination [22, 97–99] and that circulating vaccine-derived poliovirus (cVDPV) outbreaks are unknown [9, 43] outside of isolated communities with atypical immunological and social conditions [100–102].

**Figure 9.**
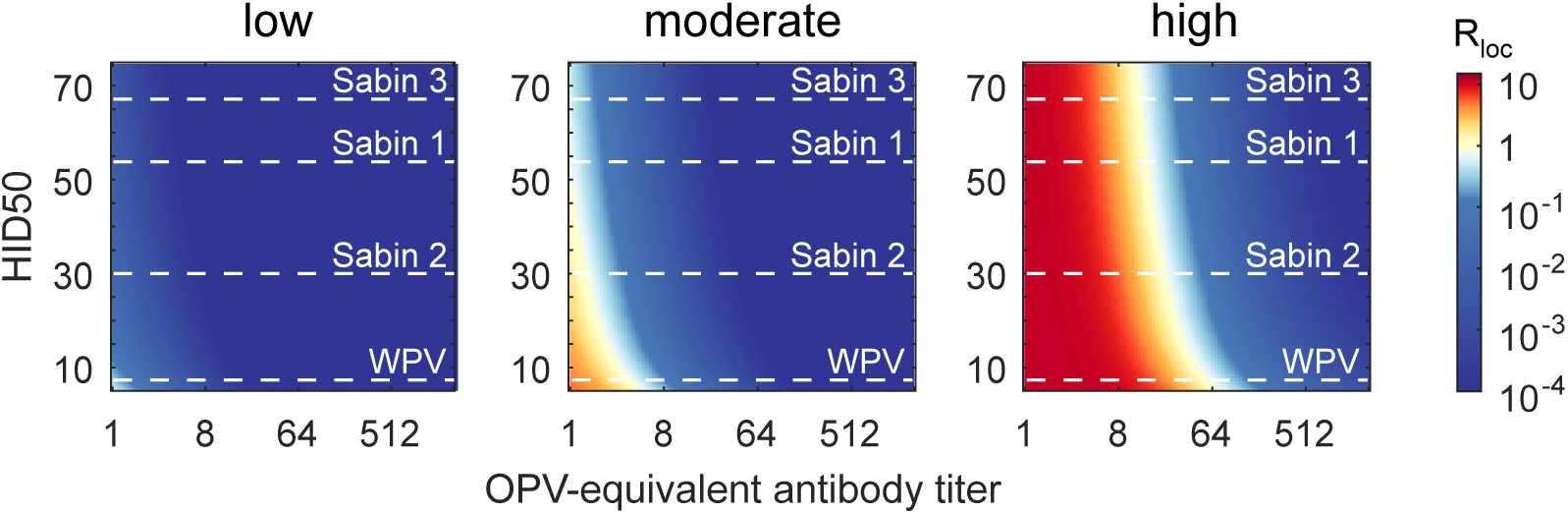
Effects of poliovirus strain on the local reproduction number. *R*_loc_ vs. the oral dose that infects 50% of immunologically-naive humans (HID50) and OPV-equivalent antibody titer for low (fecal-oral dose *T*_*ih*_ = 0.5 *µ*g/day and number of close social contacts *N*_*s*_ = 3), moderate (Houston 1960, *T*_*ih*_ = 5 *µ*g/day and *N*_*s*_ = 4), and high (UP and Bihar, *T*_*ih*_ = 230 *µ*g/day and *N*_*s*_ = 10) transmission rate settings. (Colormap, *R*_loc_; dashed lines, MLE for the HID50 of each strain (Table 1).)

In high transmission rate settings (such as UP and Bihar 2003–2008 [37]), reinfection of previously immunized people can permit community-wide epidemics if typical immunity is below a threshold level. In the example shown, one-dose OPV-equivalent immunity (*N*_Ab_ = 8) has little or no impact on *R*_loc_ for any poliovirus strain, and WPV elimination requires *N*_Ab_ *>* 64 for all. This result that WPV could persist despite *N*_Ab_ *>* 8 for most children in UP and Bihar 2003–2008 is supported by serosurveillance [103]. Prior to WPV elimination, the endemic dynamics of natural loc Loc infection and vaccination conspire to maintain typical immunity levels near 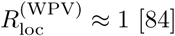, and thus the Sabin strains must have 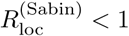, with Sabin 2 highest and Sabin 3 lowest. This result is consistent with the historical experience that vaccine-derived outbreaks have only been observed after genetic reversion has restored WPV-like properties in places where the WPV serotype has been eliminated [43, 44], and that type 2 cVDPV are most common [9]. However, if poliovirus is re-introduced after elimination into a high transmission rate setting with insufficient immunity, our model predicts that epidemic dynamics will be similar for all strains: 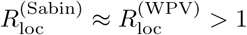 is determined by the number of social contacts and is insensitive to differences in infectiousness of the Sabin strains.

Our results above, combined with the observation that cVDPV outbreaks have only been observed at rates of roughly one per year per 250 million childen at risk under fifteen years of age [9], indicate that settings where the transmission rate for the Sabin strains is high have been rare. To evaluate how community susceptibility to Sabin 2 transmission will change due to anticipated vaccination policy changes after WPV eradication [95], we considered four scenarios for childhood immunity against type 2 poliovirus in Fig 10A. The tOPVx3 scenario describes pre-cessation populations where all index, household member, and close social contacts had achieved maximum immunity prior to waning. The bOPV & tOPVx3 scenario applies in the first two to three years after type 2 cessation, when birth spacing [104] is such that the likely index child in a family has only received bOPV (and possibly IPV) but older household members and their contacts have had tOPV. The bOPV scenario applies when two or more children in a typical household are born after type 2 cessation, and the naive scenario applies in settings where all OPV immunization has stopped. Prior and up to a few years after type 2 cessation, 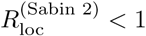 almost everywhere. However, our model predicts that 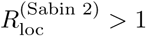 will be common when typical households have more than one child born after type 2 cessation and where hygenic practices are comparable to those of UP and Bihar in the years preceeding WPV elimination. Some moderate transmission settings may also become susceptible to Sabin 2 outbreaks once all OPV vaccination is stopped.

**Figure 10.**
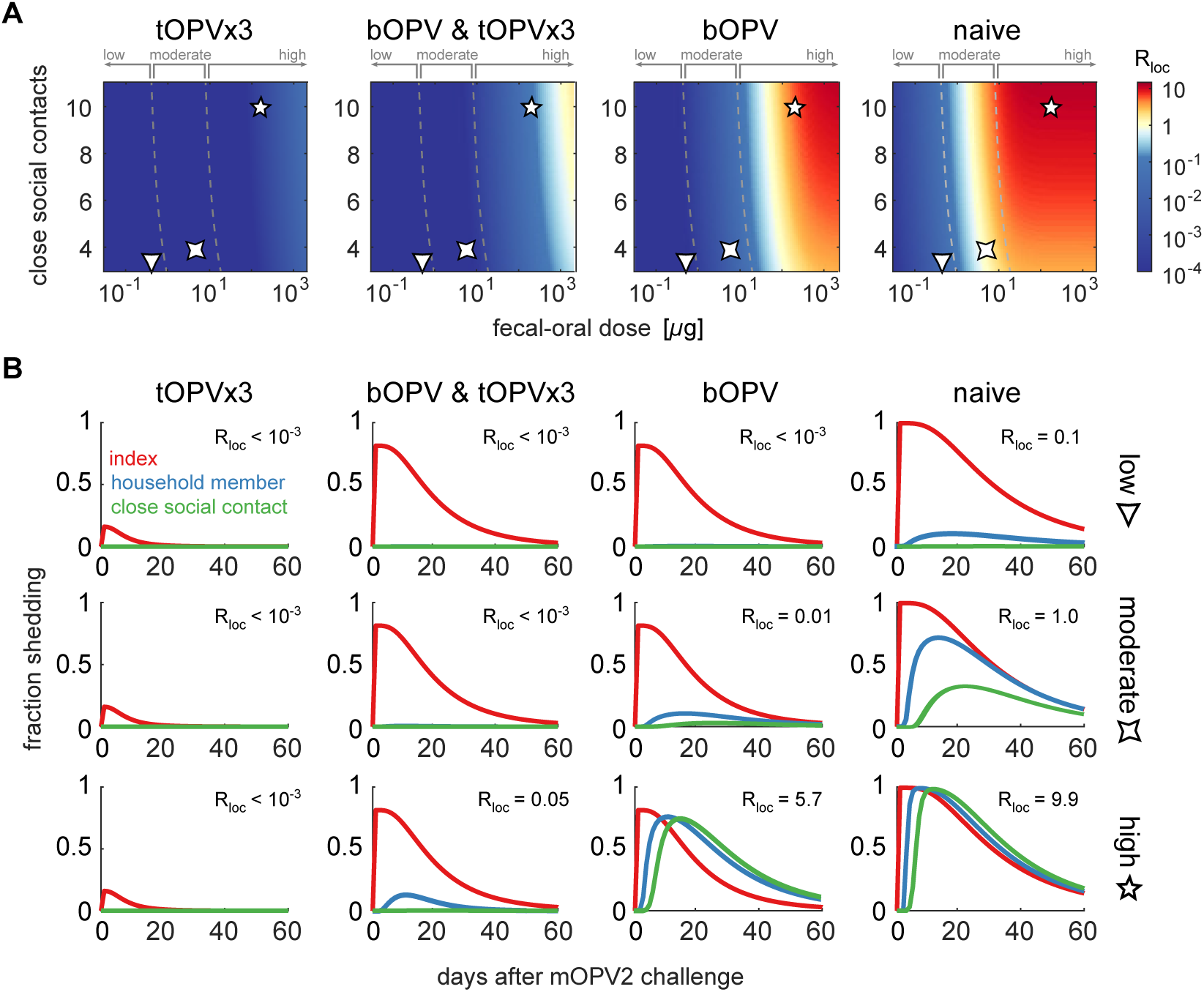
**The effects of vaccination policy on Sabin 2 transmission** for four immunity scenarios: tOPVx3 (index and household/social contact *N*_Ab_ = 512), bOPV & tOPVx3 (index *N*_Ab_ = 512 and household/social contact *N*_Ab_ = 256), bOPV (index *N*_Ab_ = 8 and household/social contact *N*_Ab_ = 2), and naive (index and household/social contact *N*_Ab_ = 1). (A) Local reproduction number vs. fecal-oral dose and number of close social contacts. (Colormap, *R*_loc_; dashed lines, transmission rate category boundaries from Fig 8B; symbols, example low, moderate, and high transmission rate settings.) (B) Maximum likelihood estimates of the fraction shedding for each subject type after mOPV2 challenge in young index children, for each immunity scenario and example transmission rate setting in panel A.

To relate local reproduction number to data that can be collected in the field, Fig 10B shows our maximum likelihood estimates for the fraction of index children, household members, and close social contacts that shed after mOPV2 challenge of the index child. In well-protected communities (*R*_loc_ ≪ 1), the model predicts little to no measurable transmission from index children infected with Sabin 2, but when *R*_loc_ ≫ 1, Sabin 2 transmission from index children to unvaccinated contacts will be nearly indistinguishable from WPV [24, 35, 37].

Fig 11 shows the sensitivity of the local reproduction number in immunologically-naive settings to social distance, measured in terms of the fecal-oral dose (*T*_*hs*_) and the household member to social contact interaction rate (*D*_*hs*_). In moderate transmission rate settings such as Houston 1960, *R*_loc_ declines rapidly with increasing social distance even in the absence of immunity. Relative to the calibrated parameters that describe transmission among close contacts, a ten-fold reduction in either fecal-oral dose or interaction rate reduces *R*_loc_ from near 1 to less than 0.1. In moderate transmission rate settings, significant transmission requires regular, undiluted contact, and so Sabin 2 is unlikely to spread outside of the communities it is delivered to. However, in high transmission rate settings such as UP and Bihar 2003–2008, *R*_loc_ can remain above 1 across two orders of magnitude in fecal-oral dose or interaction rate—and above 0.1 across three. Under these conditions, transmission does not require undiluted fecal-oral contact, and Sabin 2 can escape local communities via social interactions that take place only a few times per year.

**Figure 11.**
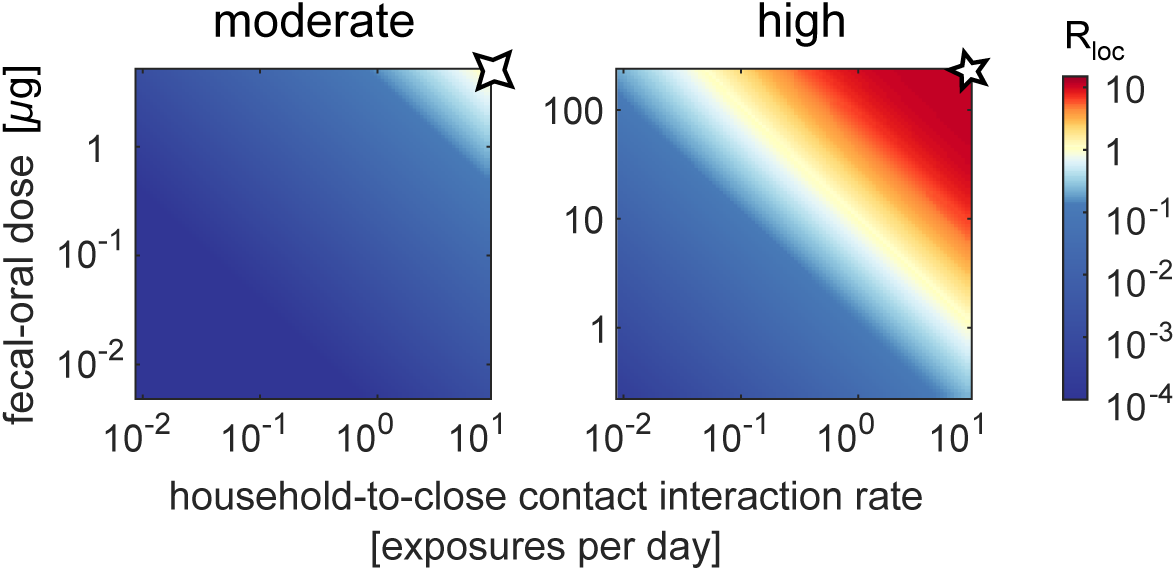
**Effects of increasing social distance on the local reproduction number of Sabin 2** in immunologically-naive populations (*N*_Ab_ = 1). Local reproduction number vs. household member to social contact interaction rate and fecal-oral dose, for moderate (*N*_*s*_ = 4, *T*_*ih*_ = 5 *µ*g/day, *Ths ≤* 5 *µ*g/day) and high (*N*_*s*_ = 10, *T*_*ih*_ = 230 *µ*g/day, *Ths ≤* 230 *µ*g/day) transmission rate settings. (Colormap, *R*_loc_; symbols, example parameter values from Fig 10.)

## Discussion

We have shown how the effects of immunity on poliovirus shedding and susceptibility to infection interact with sanitation and local interfamilial relationships to determine community susceptibility to poliovirus transmission. We found that the local reproduction number is a useful threshold statistic for characterizing the transmission rate. The highest typical levels of OPV-equivalent immunity in our model predict *R*_loc_ *<* 1 for all strains in all settings. In low and moderate transmission rate settings, we inferred that the Sabin strains have 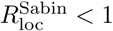 due to attenuated infectiousness relative to WPV (Fig 9), and thus significant person-to-person Sabin transmission is unlikely regardless of population immunity. Moderate transmission rate settings are at risk of outbreaks from WPV or imported (wild-like) cVDPV, but are unlikely to generate indigenous Sabin-derived outbreaks. However, in high transmission rate settings with low population immunity—a situation than can only exist in the absence of endemic transmission and OPV vaccination—our model predicts that the transmission rate of the Sabin strains if re-introduced will exceed all levels experienced prior to OPV cessation, approaching that of WPV and with highest risk for Sabin 2.

Other published mathematical models known to us have explored the effects of immunity on Sabin transmission [20, 26, 31, 32]. Despite substantial methodological differences, all are in agreement that the Sabin strains will have reproduction numbers above one in high transmission rate settings with low population immunity. In addition to novel results for dose response and waning, the key innovation of our work is its direct connection from individual-level measures of shedding and susceptibility obtained by stool surveys to assessment of community susceptibility (Fig 10). A recent application of this model to a field transmission study in Matlab, Bangladesh [86] found that moderate transmission rate conditions exist in a low-income, high-density community in the developing world where comprehensive maternal and child health care and improved sanitation systems are in place [105]. The key limitation of our model is that, while it can predict when the outbreak risk from OPV vaccination is negligible, it cannot address the absolute probability, severity, or geographic scope of outbreaks when they are possible without incorporating additional structural assumptions and calibration data about socially-distant transmission. We discuss the relevance of our results for interpreting the history of and implications for vaccination policy in the polio eradication endgame [95] below.

Before polio vaccination, most people were immunized against subsequent polio infection by natural exposure to WPV at young ages. The Sabin strains dramatically lowered the burden of paralytic disease by producing unprecedentedly high levels of immunity and displacing WPV. OPV cessation is intended to eliminate the residual disease burden caused by the Sabin strains [5, 18], but stopping OPV vaccination will reduce global immunity against poliovirus transmission to unprecedentedly low levels.

Many high-income countries with good sanitation and smaller family sizes have maintained polio elimination solely through the routine use of IPV [7, 106, 107]. Although IPV alone has little to no impact on susceptibility or shedding in stool (Fig 7), our results show that the fecal-oral route alone is incapable of supporting epidemic transmission in low transmission rate settings (Fig8). When the oral-oral route is required to permit significant transmission, our results indicate that it is possible for IPV alone to prevent outbreaks by reducing oral shedding [34, 38–41]. The Netherlands is an example of a country where IPV alone has been sufficient. In 1978 and 1992, there were outbreaks of WPV, but virus was found almost exclusively within high-risk groups who refused vaccination and no evidence of circulation in the well-vaccinated population was found [96, 108–110]. Furthermore, many countries that could not have eliminated WPV with IPV alone a few decades ago appear now to be adequately protected. The United States is an example. While there is some evidence that IPV alone could reduce WPV transmission among middle- and upper-class families in 1960 [39], IPV vaccination of subjects with no prior exposure to live poliovirus had no impact on transmission for both Sabin and WPV strains in communities with low socioeconomic status [36, 111]. However, since 2000, the United States has only used IPV [106] and yet has remained polio-free in all vaccinated populations [101] despite extensive international connections and cross-border mixing with OPV-using countries [112].

The 2013 WPV outbreak in Israel shows the limits of IPV to prevent transmission. Eight years after Israel switched from using both OPV and IPV to using IPV only, a type 1 WPV outbreak was tracked via sewage surveillance from February 2013 until April 2014 [113, 114]. Most infections were found in children born after the switch despite 93 + % coverage with two or more doses of IPV and waning immunity in older people [15, 115]. A recent model estimated that the effective reproduction number of WPV among children in the Bedouin community in which transmission was most common was 1.8 [15]; the corresponding reproduction numbers for the Sabin strains, assuming our model of infectivity, are 0.4 and below. Our interpretation is that Israel in 2013 was an example of a moderate transmission rate setting where WPV can persist despite comprehensive IPV vaccination but the Sabin strains cannot [116–118].

In the above scenarios, our model predicts that OPV cessation is stable. OPV can be used to interrupt outbreaks of WPV or imported (WPV-like) cVDPV, and the persistence of vaccine-derived strains is unlikely within (Fig 10A) or outside (Fig 11A) the outbreak response zone. However, in high transmission rate settings with low immunity, we expect that Sabin transmission to unvaccinated contacts within outbreak response regions will be common (Fig 10) and significant transmission to socially-distant contacts will occur (Fig 11B). In these settings, OPV cessation is inherently unstable—if poliovirus is re-introduced, there is no guarantee that transmission can be stopped and new cVDPV prevented without restarting OPV vaccination in all high transmission rate settings.

Our conclusion that global OPV cessation is unstable follows from the inference that doses acquired via fecal-oral exposure can be much higher in the developing world than they were in the countries where Sabin OPV was first studied and where OPV cessation has already been successful (Fig 8). The time when instability will reveal itself is uncertain. Our model predicts that two or more children per family born after cessation are required to support Sabin 2 outbreaks in most high transmission rate settings (Fig 10). The median birth spacing in most bOPV-using countries is 24–36 months [104]. Thus, we predict that between early 2018 and mid-2019, the risk of establishing type 2 cVDPV will increase substantially in many regions of the developing world that have not received post-cessation mOPV2 campaigns. The cross-immunity from bOPV against type 2 (with or without IPV) does not alter this conclusion.

Our estimate of 2 to 3 years to increased cVDPV2 risk upon Sabin 2 re-introduction is consistent with predictions from other models [20, 26, 32, 33] and is compatible with the known epidemiology of cVDPV2 outbreaks. The first known example of widespread circulation following a small release of Sabin 2 took place in Belarus in 1965, but was only confirmed as such in 2003 [119]. Two years after a local experiment in type 2 OPV cessation, tOPV given to forty children likely spread Sabin-derived poliovirus throughout a city of 160 thousand people for at least ten months. In northern Nigeria, after widespread vaccine refusal in 2003 and 2004 [120], restoration of tOPV vaccination seeded twelve independent type 2 Sabin-derived, including the largest known outbreak of cVDPV2 in history [44].

The introduction of IPV in routine immunization globally between 2014 and 2016 aimed to provide protection against poliomyelitis to children born after OPV cessation [121]. But without substantial improvements in sanitation, IPV supply [7], and routine immunization coverage, we expect that IPV alone is insufficient to protect against poliovirus circulation in all settings. In pursuit of high vaccine efficacy with low virulence [4, 122], Sabin selected strains that are 1,000–10,000 times less likely to cause paralysis than WPV [5], but only four to ten times less infectious (Table 1). In the absence of population immunity, the differences in infectiousness are insufficient to limit transmission and prevent the evolutionary restoration of virulence [43]. As a consequence, Sabin OPV will be insufficient to guarantee protection from circulation in high transmission settings [20, 123, 124]. To secure polio eradication for all times and in all conditions, we believe improved vaccines that produce infection-blocking immunity without the risks of Sabin OPV are required. Genetically-stabilized, engineered live vaccines are in development and promise the benefits of Sabin OPV without the risks [125–127], and adjuvanted IPV may provide a complementary route to a new effective vaccine [128].

Regardless of the challenges detailed above, Sabin OPV vaccination is always preferable to natural infection by WPV or cVDPV. Thus, mass vaccination with OPV remains the most effective intervention to eliminate poliovirus transmission [3], and the continued use of mOPV2 in regions experiencing type 2 outbreaks is warranted [18] despite concerns about poliovirus containment [19]. For risk mitigation, our model shows the value of healthy contact stool surveillance. The fraction of vaccine recipients and unvaccinated contacts shedding is a direct probe of population immunity and the local transmission rate, and our results provide a rubric to categorize the risk of subsequent outbreaks. Furthermore, with data about fecal-oral contamination (whether from studies of other enteric diseases or sanitation), our model can be calibrated to predict transmission rates in the absence of poliovirus and may thus have predictive value far into the post-cessation future. To go from outbreak risk categorization to risk quantification, continuing work to better understand the relationships between local and non-local transmission is needed [129–131].

## Acknowledgments

We would like to thank Benoît Raybaud and Qinghua Long (IDM) for developing the online interactive data visualization tools, Laina Mercer, Steve Kroiss (IDM) for critical feedback on the manuscript, Ananda Bandyopadhyay, John Modlin, Arend Voorman (Bill & Melinda Gates Foundation) and Hil Lyons (IDM) for useful discussions that motivated this work and critical feedback, and James S. Koopman (University of Michigan) for motivating us to examine waning immunity. We also thank Ananda Bandyopadhyay for early access to data and results from the bOPV challenge studies.

## S1 Parameter table

The values of all parameters used in the model, both from calibration and in the Results presentation, are shown in Table S1.

**Table S1.**
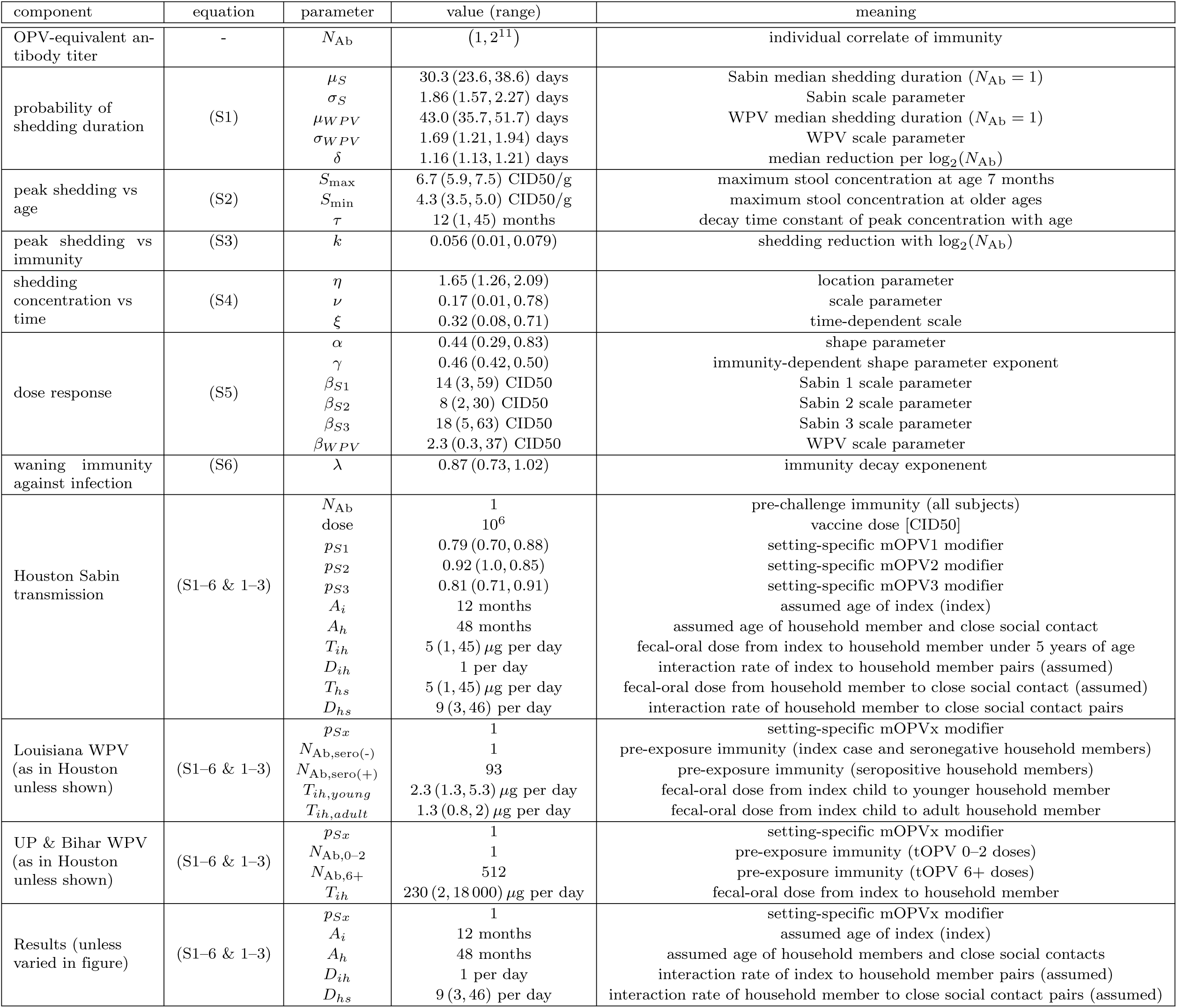
Parameter table.

| component | equation | parameter | value (range) | meaning |
| --- | --- | --- | --- | --- |
| OPV-equivalent antibody titer | - | $N_{Ab}$ | (1, 2<sup>11</sup>) | individual correlate of immunity |
| probability of shedding duration | (S1) | $\mu_S$<br>$\sigma_S$<br>$\mu_{WPV}$<br>$\sigma_{WPV}$<br>$\delta$ | 30.3 (23.6, 38.6) days<br>1.86 (1.57, 2.27) days<br>43.0 (35.7, 51.7) days<br>1.69 (1.21, 1.94) days<br>1.16 (1.13, 1.21) days | Sabin median shedding duration ($N_{Ab} = 1$)<br>Sabin scale parameter<br>WPV median shedding duration ($N_{Ab} = 1$)<br>WPV scale parameter<br>median reduction per $\log_2(N_{Ab})$ |
| peak shedding vs age | (S2) | $S_{max}$<br>$S_{min}$<br>$\tau$ | 6.7 (5.9, 7.5) CID50/g<br>4.3 (3.5, 5.0) CID50/g<br>12 (1, 45) months | maximum stool concentration at age 7 months<br>maximum stool concentration at older ages<br>decay time constant of peak concentration with age |
| peak shedding vs immunity | (S3) | $k$ | 0.056 (0.01, 0.079) | shedding reduction with $\log_2(N_{Ab})$ |
| shedding concentration vs time | (S4) | $\eta$<br>$\nu$<br>$\xi$ | 1.65 (1.26, 2.09)<br>0.17 (0.01, 0.78)<br>0.32 (0.08, 0.71) | location parameter<br>scale parameter<br>time-dependent scale |
| dose response | (S5) | $\alpha$<br>$\gamma$<br>$\beta_{S1}$<br>$\beta_{S2}$<br>$\beta_{S3}$<br>$\beta_{WPV}$ | 0.44 (0.29, 0.83)<br>0.46 (0.42, 0.50)<br>14 (3, 59) CID50<br>8 (2, 30) CID50<br>18 (5, 63) CID50<br>2.3 (0.3, 37) CID50 | shape parameter<br>immunity-dependent shape parameter exponent<br>Sabin 1 scale parameter<br>Sabin 2 scale parameter<br>Sabin 3 scale parameter<br>WPV scale parameter |
| waning immunity against infection | (S6) | $\lambda$ | 0.87 (0.73, 1.02) | immunity decay exponent |
| Houston Sabin transmission | (S1-6 & 1-3) | $N_{Ab}$<br>dose<br>$p_{S1}$<br>$p_{S2}$<br>$p_{S3}$<br>$A_i$<br>$A_h$<br>$T_{ih}$<br>$D_{ih}$<br>$T_{hs}$<br>$D_{hs}$ | 1<br>10<sup>6</sup><br>0.79 (0.70, 0.88)<br>0.92 (1.0, 0.85)<br>0.81 (0.71, 0.91)<br>12 months<br>48 months<br>5 (1, 45) $\mu\text{g}$ per day<br>1 per day<br>5 (1, 45) $\mu\text{g}$ per day<br>9 (3, 46) per day | pre-challenge immunity (all subjects)<br>vaccine dose [CID50]<br>setting-specific mOPV1 modifier<br>setting-specific mOPV2 modifier<br>setting-specific mOPV3 modifier<br>assumed age of index (index)<br>assumed age of household member and close social contact<br>fecal-oral dose from index to household member under 5 years of age<br>interaction rate of index to household member pairs (assumed)<br>fecal-oral dose from household member to close social contact (assumed)<br>interaction rate of household member to close social contact pairs |
| Louisiana WPV (as in Houston unless shown) | (S1-6 & 1-3) | $p_{Sx}$<br>$N_{Ab,sero(-)}$<br>$N_{Ab,sero(+)}$<br>$T_{ih,young}$<br>$T_{ih,adult}$ | 1<br>1<br>93<br>2.3 (1.3, 5.3) $\mu\text{g}$ per day<br>1.3 (0.8, 2) $\mu\text{g}$ per day | setting-specific mOPVx modifier<br>pre-exposure immunity (index case and seronegative household members)<br>pre-exposure immunity (seropositive household members)<br>fecal-oral dose from index child to younger household member<br>fecal-oral dose from index child to adult household member |
| UP & Bihar WPV (as in Houston unless shown) | (S1-6 & 1-3) | $p_{Sx}$<br>$N_{Ab,0-2}$<br>$N_{Ab,6+}$<br>$T_{ih}$ | 1<br>1<br>512<br>230 (2, 18 000) $\mu\text{g}$ per day | setting-specific mOPVx modifier<br>pre-exposure immunity (tOPV 0-2 doses)<br>pre-exposure immunity (tOPV 6+ doses)<br>fecal-oral dose from index to household member |
| Results (unless varied in figure) | (S1-6 & 1-3) | $p_{Sx}$<br>$A_i$<br>$A_h$<br>$D_{ih}$<br>$D_{hs}$ | 1<br>12 months<br>48 months<br>1 per day<br>9 (3, 46) per day | setting-specific mOPVx modifier<br>assumed age of index (index)<br>assumed age of household members and close social contacts<br>interaction rate of index to household member pairs (assumed)<br>interaction rate of household member to close social contact pairs (assumed) |

## S2 Within-host model

### S2.1 Sources of data on shedding and oral susceptibility to infection

Almost all relevant studies on OPV shedding, acquisition, and transmission published prior to 2012 were reviewed by Duintjer Tebbens et al [1]. Digitized data on shedding duration and concentration of poliovirus in stool were taken from the Supplementary Material of Behrend *et al* [2], corrected where discrepancies were noticed, and studies involving bOPV were added [3–5]. Dose response data were digitized from the cited references [6–9]. The analyses are broadly inclusive of published data, but this paper does not represent a systematic review with pre-specified exclusion criteria. Whole studies and trial arms were excluded if they reported evidence of substantial unmeasured exposure to poliovirus prior to OPV challenge [10–16] or when data across serotypes could not be disaggregated [17]. We included OPV challenge studies where subjects experienced low levels of natural exposure to WPV or OPV during the study, provided published evidence showed that most of the subjects were unaffected [7, 9, 18, 19]. A summary of all included data describing vaccination schedules, OPV challenge formulation or WPV exposure, ages, and shedding and dose reponse data, and possible natural exposure is provided in Table S2 [3–9, 18–31]. For a deeper discussion of data quality from reviewed studies, see Duintjer Tebbens *et al* [1].

**Table S2.** OPV challenge studies included in analysis. Ages rounded to nearest month. “Live virus exposure” indicates possible uncontrolled exposure to OPV or WPV during study. More detailed information about the included and considered but excluded studies can be found in the digitized data tables available at famulare.github.io/cessationStability/. * IPV administered at same time as OPV; † IPV administered alone but after prior OPV.

| RI schedule | RI schedule | challenge | age at challenge | location | publication date | live virus exposure | shedding duration | shedding titer | dose response | reference |
| --- | --- | --- | --- | --- | --- | --- | --- | --- | --- | --- |
| seronegative natural | - | mOPV1<br>mOPV2<br>mOPV3 | 5 y<br>20 y | Netherlands | 1959 | yes | yes | no | no | Verlinde1959 [20] |
| seronegative IPVx2 | - | mOPV1<br>mOPV2 | 13 m<br>11 m | UK | 1961 | no | yes | yes | no | Dick1961 [21] |
| seronegative | - | mOPV2 | 12 m | UK | 1961 | no | no | yes | yes | Dane1961 [32] |
| seronegative | - | mOPV1<br>tOPV | 2 y | USA | 1961 | no | yes | no | no | Horstmann1961 [22] |
| unvaccinated | - | mOPV1<br>bOPV | 0 m | USA | 1962 | no | yes | no | no | Holguin1962 [23] |
| unvaccinated<br>tOPVx3<br>IPVx3<br>IPVx4 | -<br>7,8,9 m<br>2,3,4 m<br>2,3,4,15 m | mOPV1 | 6 m<br>16 m<br>6 m<br>16 m | UK | 1966 | yes | yes | yes | yes | Henry1966 [7] |
| unvaccinated | - | mOPV1<br>mOPV2<br>mOPV3 | 2 y | Japan | 1966 | no | yes | no | no | Takatsu1966 [24] |
| unvaccinated | - | mOPV1<br>mOPV2<br>mOPV3<br>tOPV | 1 y | USA | 1967 | no | yes | no | no | Benyesh-Melnick1967 [25] |
| unvaccinated | - | mOPV1 | 2 m | UK | 1981 | no | no | no | yes | Minor1981 [8] |
| unvaccinated<br>tOPVx1 | -<br>0 m | tOPV | 0 or 2 m<br>2 m | China | 1986 | no | yes | no | no | Dong1986 [26] |
| tOPVx3<br>IPVx3 | 2,4,18 m | tOPV | 2 y | USA | 1991 | yes | yes | no | yes | Onorato1991 [9] |
| unvaccinated<br>IPVx2 | -<br>2,3 m | tOPV | 2 m<br>4 m | Romania | 1997 | yes | yes | no | no | Ion-Nedelcu1997 [18] |
| unvaccinated<br>tOPVx1<br>tOPVx2 | -<br>7 m<br>7,8 m | tOPV | 7 m<br>8 m<br>9 m | France | 1997 | no | yes | no | no | Mallet1997 [27] |
| IPVx3 | 4,6,12 m | mOPV3 | 18 m | Finland | 1999 | no | yes | yes | no | Piirainen1999 [28] |
| seronegative natural | - | mOPV1<br>mOPV3 | 65 y | Netherlands | 2005 | yes | yes | no | no | Abbink2005 [29] |
| unvaccinated<br>tOPVx2<br>IPVx2 | -<br>2,4 m<br>2,4 m | tOPV | 2 m<br>6 m<br>6 m | USA | 2005 | yes | yes | no | no | Laassri2005 [19] |
| mOPV1x1<br>tOPVx1 | 0 m | mOPV1 | 1 m<br>2 y | Egypt | 2008 | no | yes | no | no | El-Sayed2008 [30] |
| IPVx2<br>& tOPVx2<br>IPVx3<br>& tOPVx3 | 2*,4*,7 m<br>2*,4*,7,13*<br>m | tOPV | 10 m<br>16 m | Israel | 2008 | no | yes | no | no | Swartz2008 [31] |
| tOPVxN<br>IPV boost | campaigns | mOPV1<br>mOPV3 | 1,5,10 y | India | 2014 | yes | no | no | yes | Jafari2014 [3] |
| IPVx1<br>& bOPVx2<br>IPVx2<br>& bOPVx1<br>IPVx3 | 2*,3,4 m<br>2*,3*,4 m<br>2*,3*,4* m | mOPV2 | 6 m | Chile | 2015 | no | yes | yes | no | O’Ryan2015 [4] |
| bOPVx3<br>& IPVx1<br>bOPVx3<br>& IPVx2<br>bOPVx3<br>tOPVx3 | 2,3,4* m<br>2,3,4*,8†<br>m<br>2,3,4 m<br>2,3,4 m | mOPV2 | 5 or 9 m<br>9 m<br>5 m<br>5 m | Latin America | 2016 | no | yes | yes | no | Asturias2016 [5] |

### S2.2 Shedding duration after OPV challenge or WPV infection

We assumed a log-normal survival distribution for the shedding duration given infection:

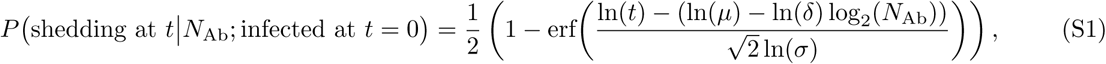

where *N*_Ab_ is the OPV-equivalent antibody titer at *t* = 0, *µ* is the median duration in days for immunologically-naive individuals (*N*_Ab_ = 1), *δ* describes the decrease in median duration with increasing immunity, and *σ* describes the shape of the distribution. The median durations and OPV-equivalent antibody titers shown in Fig. 2 were estimated under this model. Figure S1 shows the model maximum likelihood estimates (MLE) and 95% confidence intervals (CI) for the shedding duration distribution at low and high OPV-equivalent antibody-titers. An earlier version of this model was published within the supplemental software of Behrend *et al* [2] but was not described in that paper, and the model was used without derivation in references [33, 34]. Given the approximate aggregated survival distributions in Fig 1A, we estimated approximate maximum likelihood parameters of the shedding duration model using binomial maximum likelihood (assuming independent samples). We used parametric bootstrap to estimate confidence intervals.

**Figure S1.**
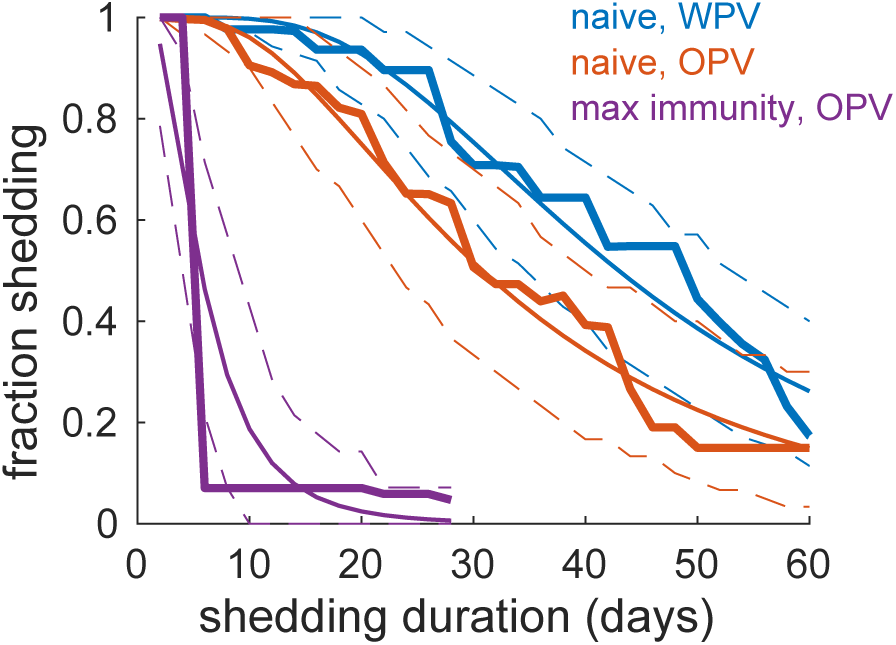
Shedding duration probability for immunologically-naive and maximally-immune individuals. Empirical shedding duration reverse-cumulative distributions, model maximum likelihood estimate, and 95% confidence interval shown.

We estimated that the WPV shedding duration in immunologically-naive children was 43.0 (35.7, 51.7) days from longitudinal surveillance studies of WPV incidence, significantly longer than our estimate for shedding duration after OPV challenge, (30.3 (23.6, 38.6) days). To confirm that this estimate is not an artifact of differences between OPV challenge and WPV surveillance study design, we examined alternative data for the time from infection to paralysis and for shedding duration after the onset of paralysis. Casey *et al* measured that the mean time to paralysis from WPV infection is 17 days [35] and Grassly *et al* showed that the mean shedding duration after paralysis from WPV infection in UP & Bihar is 31 days [36]. The sum, 48 days, is consistent with our previous estimate.

### S2.3 Concentration of poliovirus in stool

For each trial arm that informed our concentration model [4–6, 20, 21, 28, 29], we estimated the OPV-equivalent antibody titer from the shedding duration distributions of each trial arm as above. To model the age-dependence of the concentration of poliovirus in stool, we fit an exponential model to the peak shedding concentration:

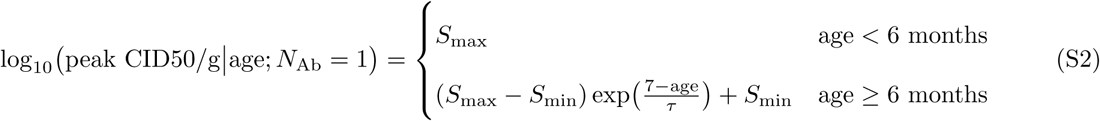

with maximum concentration *S*_max_ = 6.7 (5.9, 7.5), minumum concentration *S*_min_ = 4.3 (3.5, 5.0) CID50 per gram, and time constant *τ* = 12 (1, 45) months. We modeled the effect of pre-challenge immunity on concentration as:

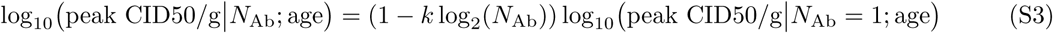

with *k* = 0.056 (0.01, 0.079). The poliovirus concentration timeseries peaks shortly after acquiring infection and declines slowly thereafter. To model viral load over time, following refs. [2, 33], we fit a quasi-log-normal shedding profile to the age-adjusted aggregated data for immunologically-naive individuals:

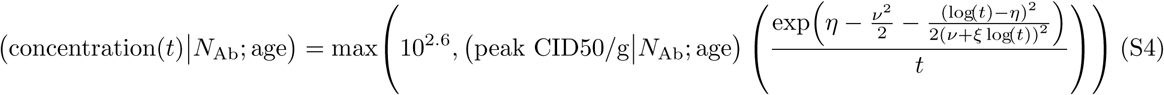

with *η* = 1.65 (1.26, 2.09), *ν* = 0.17 (0.01, 0.78), *ξ* = 0.32 (0.08, 0.71), and lower bound 10^2.6^ CID50/g to reflect the minimum reported detectable shedding.

### S2.4 Oral susceptibility to infection from OPV challenge

For each trial arm that informed our dose response model [4–9], we estimated the OPV-equivalent antibody titer from the shedding duration distributions of each trial arm as above. In order to summarize data for all doses and OPV-equivalent antibody titers, we fit a beta-Poisson dose response model for the fraction shedding after receiving an oral poliovirus dose. The beta-Poisson model is based on the assumptions that a single infectious unit (measured in CID50–the amount of poliovirus required to induce a cytopathic effect in 50% of inoculated cell or tissue culture plates) is sufficient to start an infection, that multiple infectious units contribute independently to the total probability of infection, and that the probability an infectious unit survives from initial oral exposure to the site of infection is beta-distributed [37]. Since the model in Behrend *et al* [2] fitted poorly at low doses and high immunity, we explored various parameterizations of the model and found that a parsimonious description of all the OPV challenge data was provided by:

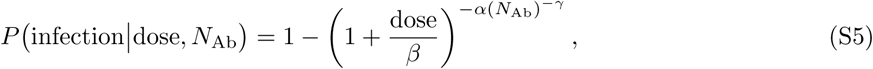

where α and *β* are the standard beta-Poisson parameters, *N*_Ab_ the OPV-equivalent antibody titer, and γ captures the reduction in shedding probability with increasing immunity.

We used the fitted dose response model to estimate the OPV-equivalent antibody titer after IPV boosting on children with many prior doses of tOPV in India [3]. The maximum likelihood estimate of the OPV-equivalent antibody titer was *N*_Ab_ = 3700 (1700, 7700) and not significantly different from the maximal immunity produced by tOPVx3 prior to any waning [5] (*N*_Ab_ = 2048 (430, 9600)).

### S2.5 Waning immunity against infection

For each trial arm that informed our waning model [3, 5, 9, 20, 29], we estimated the OPV-equivalent antibody titer from the shedding duration distributions of each trial arm as above.

The time interval between last immunization and mOPV challenge was either reported or estimated as follows. For individuals from tOPVx3 vaccine trials, intervals between last immunization and mOPV challenge ranged from 1 month [5] to 6 months [9]. To assess waning of tOPV-based immunity in older children, one study in Uttar Pradesh compared mOPV vaccine take rates in children 1, 5, or 10 years of age [3] who had previously recieved an unknown but high number of tOPV doses. To estimate the likely interval between last immunization and challenge, we assumed that children are offered up to 5 doses in the first year of life (3 RI plus 5 campaigns at 60% coverage), corresponding to roughly 2.5 months on average between last vaccination and mOPV challenge at 1 year of age. We assumed campaigns delivered 3 doses per year in ages two through four, corresponding to roughly 4 months between last vaccination and challenge at 5 years of age, and no doses after 5 years of age, corresponding to 5 years since last vaccination and challenge at 10 years of age. For this study, OPV-equivalent immunity was inferred via vaccine take rates using equation (5). Data on adult shedding after natural immunity were taken from studies in the Netherlands. From the study by Verlinde *et al* [20] in 1959, the average seropositive subject in the study was 20 years of age, and we assumed that their last infection was 5 years earlier at 15 years of age when maximum seropositivity was first achieved in the population. From the study by Abbink *et al* [29] from 2005 that measured shedding in elderly individuals upon mOPV challenge, we assumed last exposure was 45 years earlier in 1960, at roughly the year in which widespread endemic transmission ceased in the Netherlands. We included data for both seropositive and seronegative adults from the Abbink *et al* study because seronegative adults showed evidence of memory immunity and reduced shedding durations in comparison to immunologically-naive children.

We fit a power law waning model [38] to the OPV-equivalent antibody titers,

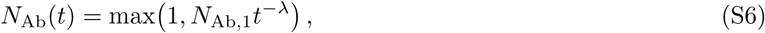

where *t* is measured in months between last immunization and oral challenge, *N*_Ab,1_ is the baseline immunity one month post-immunization, and the exponent is *λ* = 0.87 (0.73, 1.02).

## S3 Transmission model

### S3.1 Model fit

Figure S2 is an extension of Figure 6 that shows maximum likelihood estimates of the fraction shedding (prevalence) and cumulative incidence in our model for the three calibration targets. For parameters, see Table S1.

**Figure S2.**
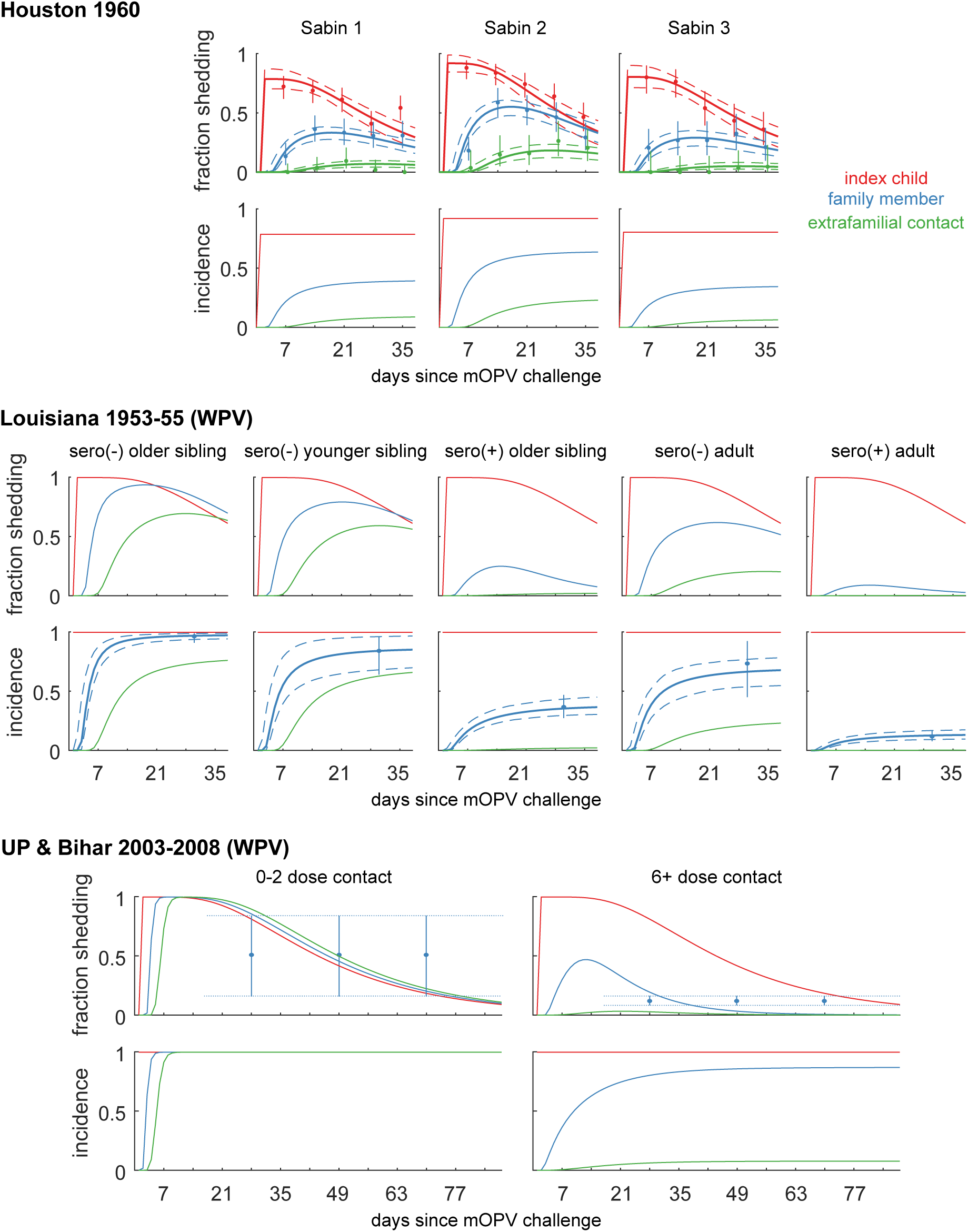
Model of fraction shedding and incidence for each calibration target. Maximum likelhood estimates (solid lines) are shown for each subject type (color) after mOPV in Houston or WPV in Lousiana and UP & Bihar. Dots-and-whiskers show calibration targets, and model 95% CI are shown for comparison. For Houston, we compared fraction shedding in stool to model prevalence. For Louisiana, cumulative incidence one month after index child infection. For UP & Bihar, we compared the mean fraction of close (direct personal) contacts shedding after index child paralysis (model prevalence and calibration target (mean over time) shown).

**Figure S3.**
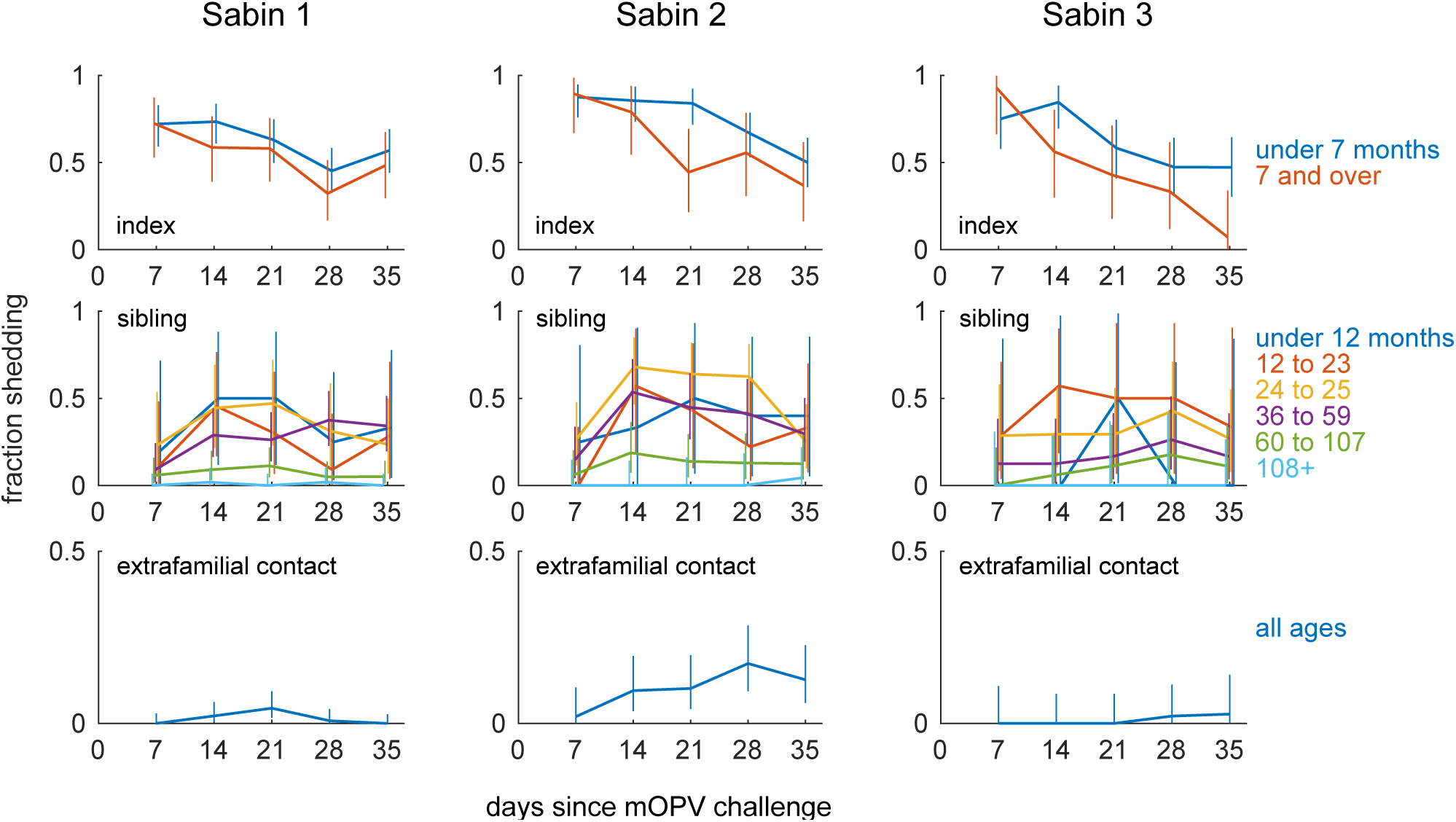
Fraction shedding by cohort and age range as originally reported. Observed fraction shedding and estimated 95% binomial confidence interval for each serotype, subject type, and reported age cohort.

### S3.2 Houston 1960

No breakdown by age was presented by Benyesh-Melnick *et al* [25] for the extrafamilial contacts of the siblings. However, because the contacts are demographically similar to the siblings and age is a significant factor for poliovirus acquisition via transmission in this setting, we used age-adjusted shedding rates in this paper. To estimate the shedding fraction in the age under 5 contact cohort, we adjusted the total reported shedding counts for each serotype as follows:

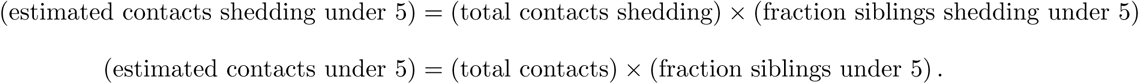

The estimated counts were rounded to the nearest integer and confidence intervals presented are based on the rounded estimated counts.

Figure S3 shows more information about the age-dependence of shedding after mOPV challenge. Older index children shed slightly less after mOPV challenge than younger children for types 2 and 3 (type 1 *p* = 0.105; type 2 *p* = 0.016; type 3 *p* = 0.025; two-tailed Fisher’s exact test). This observation was not explored in the original paper, and we propose two possible explanations. As described in Table 1 of Benyesh-Melnick *et al* [25], older index children were more likely to have received at least one dose of IPV. However, it should be noted that the original authors reported that they found no significant differences between IPV and unvaccinated index subjects, as is compatible with our metastudy. A second possibility is that stool concentrations of poliovirus are higher in young index cases, and so stool culture may be more sensitive to shedding in younger children (eq. (S2)). In the household member cohorts, there were no statistically significant differences in shedding among the age groups under 12 months, 12 to 23 months, 24 to 35 months, or 36 to 59 months for any serotype. However, there was significantly less shedding in the 60 to 107 month age group relative to the 36 to 59 age group (*p* < 0.001 for all serotypes). As stated in the main text, shedding in siblings age 60 to 107 months (5 to 9 years) is significantly below that of ages less than 5 years for all serotypes (type 1 *p* < 0.001; type 2 *p* < 0.001; type 3 *p* = 0.002). Shedding rates were very low in parents and children age 10 years and older (*<* 2%) [25], and so it is likely the transmission was direct from index child to sibling and was not mediated by infected caretakers. Shedding due to transmission-acquired type 2 was significantly more common than for types 1 and 3, and shedding due to transmission was similar for types 1 and 3 (mean prevalence: type 1 vs type 2 *p* = 0.002; type 1 vs type 3 *p* = 0.33). Primary extrafamilial contacts of siblings exhibited a similar pattern of increased type 2 shedding and comparable type 1 and 3 shedding (type 1 vs type 2 *p <* 0.001; type 1 vs type 3 *p* = 0.73). Although the authors did not describe the relationships between siblings and extrafamilial contacts in detail, it is likely that the contacts were close friends of the siblings and were directly infected by the siblings, as the authors also describe a smaller set of more socially-distant “secondary extrafamilial contacts” who “were drawn from the neighborhoods or schools attended by the siblings” and who were infected at lower rates than the primary contacts [25].

Little information about shedding in secondary contacts was provided, except to note that, summed across all trial arms, 15 of 280 secondary contacts were positive for Sabin 2 and the highest incidence rate was 13% in the secondary extra-familial contacts of tOPV recipients. Assuming the number of secondary contacts is proportional to the number of primary contacts for each trial arm, *n* = 10 of the type 2 positives were in secondary contacts of tOPV recipients (13% of trial arm total) and *n* = 5 were in secondary contacts of mOPV2 recipients (8.5% of trial arm total).

### S3.3 Louisiana 1953–1955

We calibrated model incidence to the seroconversion data reported in Gelfand *et al* [39]. Stool collection data was also available, but it reported lower levels of incidence. This was likely due to missing infections: the average interval between samples was 27 days while the average shedding duration in seropositive subjects with median *N*_Ab_ = 93 is only 16 days under our model. We assumed 100% incidence of immunologically-naive index children after WPV exposure, based on the study design that reported household member incidence conditional on detection of the child’s first natural infection with poliovirus.

### S3.4 Uttar Pradesh and Bihar 2003–2008

We calibrated our model to the estimates of mean stool prevalence after the onset of paralysis in close contacts of WPV cases reported by Grassly *et al* [36]. We assumed that the contact data best corresponded to household members in our model. Quote:

> *During identification of healthy contacts, an effort was made to include those children with the closest contact to the individual with suspected poliomyelitis, such as siblings, playmates, or residents of the same household.* [36]

To shift our model from prevalence after infection to prevalence after paralysis, we convolved our prevalence timeseries with the time-to-paralysis distribution:

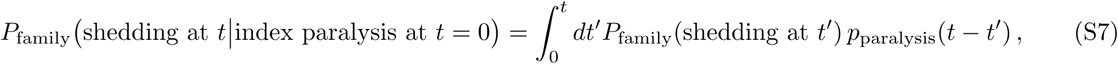

where *p*_paralysis_(*t*) was given by the histogram in Figure 2 of Casey *et al* [35]; the mean time from infection to the onset of paralysis was 17 days. The model was calibrated against the mean of eq. (S7) over the first 90 days after index child paralysis.

## Abbreviations

WPV: wild poliovirus
OPV: oral polio vaccine
tOPV: trivalent OPV
mOPV: monovalent OPV
bOPV: bivalent type 1 and 3 OPV
cVDPV: circulating vaccine-derived poliovirus
IPV: inactivated polio vaccine
CID50: the culture infectious dose that induces a cytopathic effect in 50% of infected cell or tissue cultures
HID50: the dose (measured in CID50) that infects 50% of orally-exposed and immunologically-naive humans
MLE: maximum likelihood estimate
CI: confidence inteval
UP: Uttar Pradesh

